# DNA sequence-dependent formation of heterochromatin nanodomains

**DOI:** 10.1101/2020.12.20.423673

**Authors:** Graeme J. Thorn, Christopher T. Clarkson, Anne Rademacher, Hulkar Mamayusupova, Gunnar Schotta, Karsten Rippe, Vladimir B. Teif

## Abstract

The mammalian epigenome contains thousands of heterochromatin nanodomains (HNDs) marked by di- and trimethylation of histone H3 at lysine 9, which have a typical size of 3-10 nucleosomes. However, the (epi)genetic determinants of their location and boundaries are only partly understood. Here, we compare four HND types in mouse embryonic stem cells, that are defined by histone methylases SUV39H1/2 or GLP, transcription factor ADNP or chromatin remodeller ATRX. Based on a novel chromatin hierarchical lattice framework termed ChromHL, we are able to predict HND maps with singe-nucleotide resolution. We find that HND nucleation can be rationalized by DNA sequence specific protein binding to PAX3/9, ADNP and LINE1 repeats. Depending on type of microdomains, boundaries are determined either by CTCF binding sites or by nucleosome-nucleosome and nucleosome-HP1 interactions. Our new framework allows predicting how patterns of H3K9me2/3 and other chromatin nanodomains are established and changed in processes such as cell differentiation.

---

Cell type specific gene expression programs are established by distinct patterns of active and silenced chromatin states. One important type of a repressive heterochromatin state is characterized by di- or trimethylation of histone H3 lysine K9 (H3K9me2/3) and has heterochromatin protein 1 (HP1) as a marker^1,2^. Heterochromatin can be ectopically induced by tethering HP1 or enzymes responsible for H3K9 methylation such as SUV39H1 (KMT1A) and SUV39H2 (KMT1B) to chromatin^3–5^. Remarkably, the size of domains found in the mammalian epigenome that carry the H3K9me2/3 mark, ranges from a few hundred to millions of base pairs (bp). A number of studies have investigated large heterochromatin domains in relation to genome architecture and function^6^ and several theoretical models have been introduced to describe the underlying molecular mechanisms^4,5,7–17^. These models typically include DNA-protein binding and enzymatic reactions to account for epigenetic phenomena or the mechanism to establish bistable states. Spreading of a given modification to adjacent nucleosomes on the chain is explained by nearest-neighbour feedback mechanisms^18^ as well as long-range interactions^19^, e.g. through looping of the nucleosome chain. Several mechanisms have been proposed to explain what stops heterochromatin spreading and sets domain boundaries. On the one hand, the dynamic properties of the nucleosome chain can inherently limit the interactions of a nucleosome that could propagate H3K9me3 modifications in the presence of counteracting enzymatic activities that remove this mark^13,20^. In addition, an island of nucleosomes marked by phosphorylation of histone H3 at serine 10^21^, nucleosome-depleted regions^22^ or DNA-bound molecules such as RNA polymerase or CTCF^23–26^ could act as boundary elements that interfere with the nearest-neighbour type spreading of histone modifications. While the various models have provided a wealth of insight, they are not well suited to rationalize the genome-wide formation of H3K9me2/me3 nanodomains. These endogenous patterns of tens of thousands of heterochromatin loci with a typical size of 0.7-2 kb are abundantly present throughout the mammalian genome. We call these regions heterochromatin nanodomains (HNDs). Their extension of around 3-10 nucleosomes is similar to the domain size determined by Micro-C^27^, corresponding to approximate dimensions of 40-70 nm^20^. Note that HNDs should not be confused with the significantly larger regions of ~200-300 kb observed microscopically that have been described recently as “chromatin nanodomains”^28^. In the present study we distinguish four different types of HNDs in mouse embryonic stem cells (ESCs). These HNDs are characterized by the enrichment of H3K9me2/3 and driven by the following factors: (i) histone methyltransferases SUV39H1 and SUV39H2 referred to here as SUV39H that set H3K9me3 marks^29,30^; (ii) methyltransferase GLP (G9a like protein, KMT1D) that catalyses the formation of H3K9me2^31^; (iii) transcription factor ADNP that recruits the chromatin remodeller CHD4 as well as HP1 for H3K9me3 mediated gene silencing^32^; (iv) chromatin remodeller ATRX that induces formation of H3K9me3 HNDs at repeat sequences^33^.

The description of the distribution of HNDs requires a DNA sequence-specific model with single nucleotide resolution applicable to the analysis of the complete mouse or human genome. This currently unmet need is addressed here by introducing the Chromatin Hierarchical Lattice (ChromHL) framework. ChromHL uses statistical mechanical lattice binding approaches^9,34,35^. It integrates DNA-sequence dependent transcription factor (TF) binding at single nucleotide resolution with larger-scale calculations of binding of proteins to nucleosomes in dependence of their histone modifications. This approach allows us to describe HND formation as a general mechanism involving DNA sequence-specific binding of nucleation factors and formation of HND boundaries that are determined mainly either by the DNA sequence or by nucleosome-nucleosome/HP1 interactions. Thus, our novel ChromHL framework identifies crucial DNA sequence and chromatin features and rationalises how distinct patterns of HNDs are established throughout the genome.

## Results

### Four types of endogenous HNDs are distinguished in ESCs

We first compared the structure and composition of four types of HNDs marked by H3K9me2/3 in ESCs. Previously published data sets were used for HNDs associated with SUV39H^29,30^, GLP^31^ and ADNP^32^. In addition, a dataset for ATRX-dependent H3K9me3 HNDs was newly generated here by ChIP-seq in wild-type (WT) ESCs and *Atrx* knock out (KO) cells (**Supplemental Fig. S1**). For SUV39H, GLP and ATRX HNDs, we called H3K9me2/3 ChIP-seq peaks separately in WT and KO conditions, and then identified peaks that were present in WT but lacking in the KO cells. This yielded 36,764 (*Suv39h1/Suv39h2* KO, H3K9me3), 48,881 (*Glp* KO, H3K9me2) and 13,113 (*Atrx* KO, H3K9me3) regions that change their H3K9 methylation state upon the knockout of the indicated protein factor in ESCs. In the case of the ADNP dataset, 4,673 H3K9me3 domains were called by intersecting H3K9me3 domains with regions bound by ADNP in wild type ESCs. Next, we calculated average profiles of HP1α, CTCF, nucleosome density and H3K9 methylation as a function of the distance from the centres of SUV39H-, GLP-, ADNP- and ATRX-associated HNDs (**Fig. 1**). In addition, the corresponding profiles for H3K27me3, H3K4me1, CpG methylation, different chromatin states defined by combinatorial histone marks from the analysis with ChromHMM and the read mappability were computed (**Supplemental Fig. S2, S3**). In all cases, HP1 binding and H3K9me3 (or H3K9me2 in the case of GLP-dependent regions) was enriched at the peak centre. In SUV39H-, GLP- and ATRX-dependent nanodomains, CTCF was found to be depleted while the nucleosome density was increased. In contrast, CTCF was significantly enriched in ADNP-associated HNDs that also showed a slight nucleosome density reduction. Interestingly, ADNP-associated HNDs overlapped with enhancers and had characteristic patterns of active histone marks such as H3K27ac, H3K4me1 and H3K36me3. Thus, this type of H3K9me3 nanodomains is likely to have functional roles different from that of the canonical silenced heterochromatin state that lacks these active histone marks. Likewise, DNA methylation was strongly enriched in SUV39H-dependent regions but not in other three types of HNDs. An interesting feature of GLP-dependent heterochromatin was a large number of repetitive regions as apparent from the drop in the mappability index. Thus, all four types of nanodomains studied here have distinct chromatin features.

**Fig. 1.**
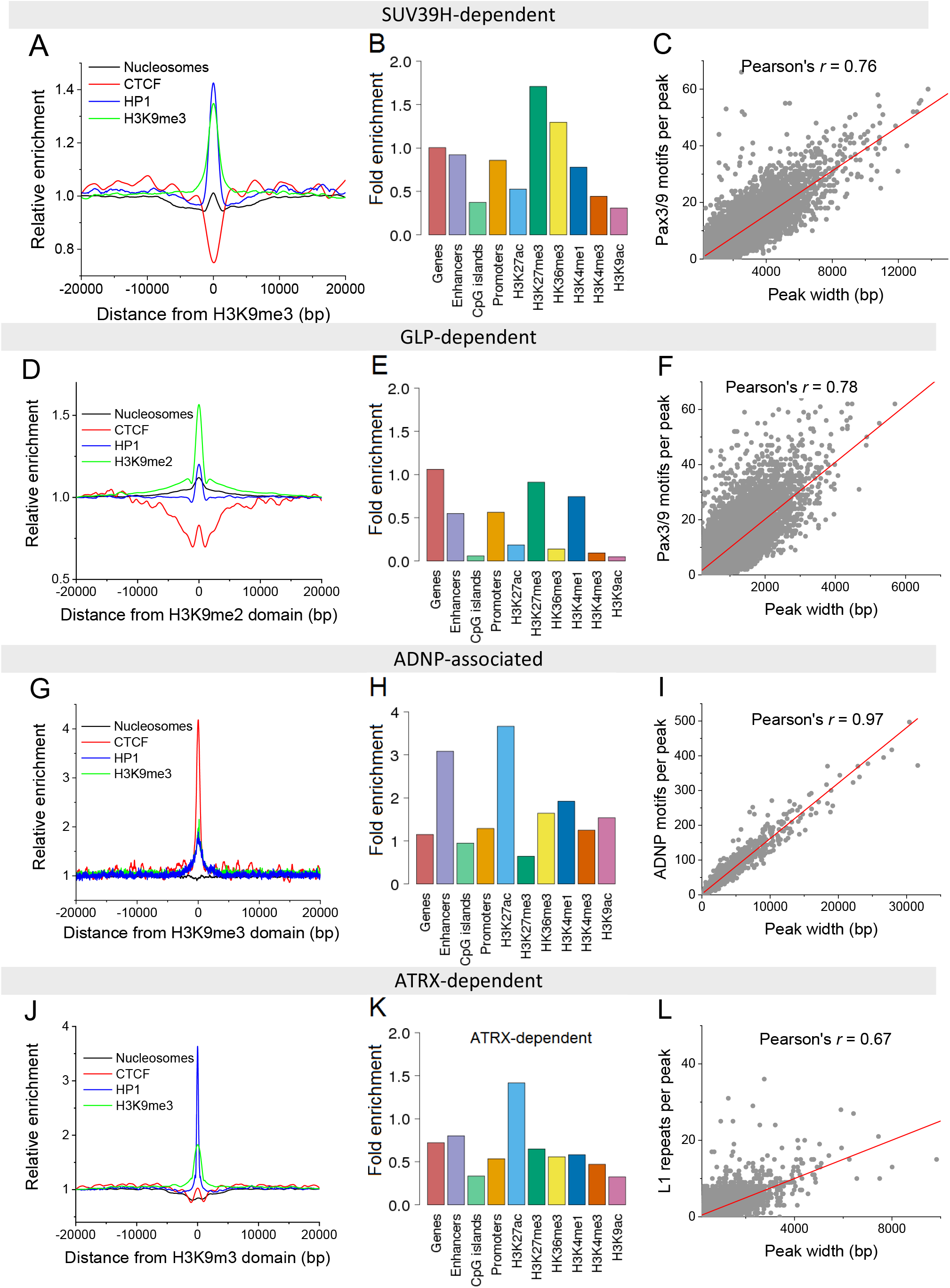
Average profiles and enrichments of chromatin features across four different endogenous HND types. Left, nucleosome occupancy, CTCF, HP1 and H3K9me3 (or H3K9me2 for GLP-dependent peaks) density. Middle, relative enrichment of different features. Right, correlation between the peak widths and the number of heterochromatininitiating motifs per peak. (**A**) SUV39H-dependent HNDs (*n* = 36,764). (**B**) GLP-dependent HNDs (*n* = 48,881). (**C**) ADNP-associated HNDs (*n* = 4,673). (**D**) ATRX-dependent HNDs (*n* = 13,113).

### Recurring sequence motifs can act as HND nucleation sites

For SUV39H-dependent H3K9me3 nanodomains, we followed the hypothesis that heterochromatin nucleation is induced by binding sites of the transcription factors PAX3 and PAX9^29^. An analysis of the SUV39H-dependent HNDs revealed that 92.4% of these indeed carried the sequence motif of the PAX3/9 binding site. The same motif was also detected in 95.9% of the GLP-dependent nanodomains, suggesting that it could also drive the formation of this nanodomain type. Interestingly, the sizes of both SUV39H- and GLP-dependent HNDs correlated well with the number of PAX3/9 motifs per corresponding HND (Pearson’s r = 0.76 and 0.78, correspondingly; **Fig. 1C, F**). For ADNP, we derived the position weight matrix (PWM) from the ChIP-seq data^32^ and used it to correlate domain extension with the number of ADNP binding motifs. This resulted in Pearson’s *r* = 0.97 for the HNDs defined from the intersection of H3K9me3 and ANDP ChIP-seq peaks (**Fig. 1I**), which means that the number of ADNP motifs per HND is an extremely good predictor of the HND size. In the case of heterochromatin formation by ATRX, the nucleation involves the recruitment of SETDB1 and/or SUV39H1 that set the H3K9me3 modification but different targeting mechanisms for these enzymes have been proposed^36–39^. Accordingly, we evaluated DNA sequence motifs that could act as nucleation sites at the 13,113 ATRX-dependent H3K9me3 nanodomains that we identified. (i) Only 4,851 (37%) of these regions contained PAX3/9 motifs. (ii) The telomeric repeat-containing RNA (TERRA) has been reported to compete with ATRX binding at the telomeric repeat sequence TTAGGG interspersed in the genome^38^. We found this sequence in 5,599 ATRX HNDs (43%). (iii) ATRX is known to bind to G-quadruplexes^39^, therefore we searched for G-quadruplex motifs^40^ within HNDs. However, only 1,026 ATRX HNDs (8%) contained such motifs (**Supplemental Fig. S7C**). (iv) IAP repeats containing a 160 bp sequence motif termed SHIN sequences have been previously reported to initiate ATRX heterochromatin^36^. However, the SHIN sequence was absent in ATRX-dependent nanodomains and <1% of ATRX HNDs intersected with annotated IAP repeats based on UCSC RepeatMasker. (v) We analysed other repeat sequences and found that all ATRX-dependent regions contained at least one LINE1 repeat (Table S1). Pearson’s correlation between the size of ATRX-dependent HNDs and the number of L1 occurrences per HND reached r = 0.67 (Fig. 1L). Thus, for all four different types of HND we identified a DNA sequence motif that could act as nucleation site (**Table 1**).

**Table 1.** Chromatin features of different endogenous HND types

|  | **SUV39H<sup>a</sup>** | **GLP<sup>a</sup>** | **ADNP<sup>a</sup>** | **ATRX<sup>a</sup>** |
| --- | --- | --- | --- | --- |
| **Marker modification** | H3K9me3 | H3K9me2 | H3K9me3 | H3K9me3 |
| **HND number** | 36,764 | 48,881 | 4,673 | 13,113 |
| **Typical HND size (kb)<sup>b</sup>** | 2.0 | 0.9 | 1.1 | 0.7 |
| **Nucleation motif** | PAX3/9 | PAX3/9 | ADNP | LINE1 subtype |
| **Correlation of HND size and nucleation motif<sup>c</sup>** | 0.76 | 0.78 | 0.97 | 0.67 |
| **HNDs with motif (%)** | 92.4 | 95.9 | 100 | 100 |
| **CTCF contribution to domain boundaries** | yes | yes | yes | no |
| **NRL** | 189±1 bp | 189±1 bp | 175±1 bp | 189±1 bp |

<sup>a</sup> Protein factor upon which the HNDs are dependent in ESCs
<sup>b</sup> Domain size as described by the full width at half maximum of the aggregated H3K9me2/3 density plots shown in Fig. 3A-D
<sup>c</sup> Correlation coefficient between nanodomain size and the number of initiation motifs per domain.

### ChromHL predicts HNDs from a DNA sequence-informed hierarchical lattice model

We developed a DNA sequence-informed hierarchical lattice model for HND formation, called ChromHL, which allows rationalising formation of different HND types (**Fig. 2**). The ChromHL framework starts with a DNA-binding lattice model at single-nucleotide resolution to determine the arrangement of nanodomain-initiating or -limiting proteins, such as PAX3/9, ADNP and CTCF. The next hierarchy level of ChromHL is defined by a lattice model with nucleosome-size units. It describes histone modifications and binding of additional proteins such as HP1 to nucleosomes and also includes nucleosome-nucleosome interactions depending on the nucleosome state. The maps of chromatin nanodomains and bound proteins are then calculated with the transfer matrix formalism of statistical mechanics (Supplementary Materials). The input for the calculations are the DNA sequence, protein-DNA affinity matrices of size 4 x *m* for each protein type *g* which cover *m*(*g*) bp upon binding. The binding is characterised by sequence-specific binding constants *K*(*n, g*) for each genomic location *n*, cooperativity parameters *w*(*g*_1_, *g*_2_) between protein types *g*_1_ and *g*_2_, as well as protein concentrations *c*(*g*) (**Fig. 2A**). At single nucleotide resolution, we use a transfer matrix model that accounts for partial unwrapping of DNA from the nucleosome core as well as competitive, cooperative DNA binding of different protein species as detailed previously^41,42^. At the nucleosome-level, we have extended the matrix model to allow different nucleosome states *e*(n) of the lattice unit *n* in addition to the bound/unbound states defined in the basic model. In the present study, three states are considered in the model (**Fig. 2B, C**): (i) a nucleosome with unmethylated H3K9 tails, (ii) a nucleosome with methylated H3K9 tails, and (iii) CTCF is bound while a nucleosome is missing from the lattice unit. This model can be extended to include other nucleosome states as needed. The critical feature of the model is the nucleosome-nucleosome interaction potential σ(*e*_1_, *e*_2_), which depends on the states of the interacting nucleosomes, *e*_1_=1 and *e*_2_=2. Thus, if neighbouring clutches of nucleosomes belong to different states, a difference in nucleosome-nucleosome interactions between these two states can create “surface tension” at the boundary with additional energy costs. Another important feature of the model is the stabilising role on heterochromatin domain formation of nucleosome-binding proteins such as HP1. HP1 binds stronger to heterochromatin states, thus shifting the thermodynamic equilibrium towards heterochromatin formation for the regions where it is bound. Neighbouring HP1 molecules bind cooperatively, characterised by parameter w, which contributes to heterochromatin spreading beyond nucleation motifs.

**Fig. 2.**
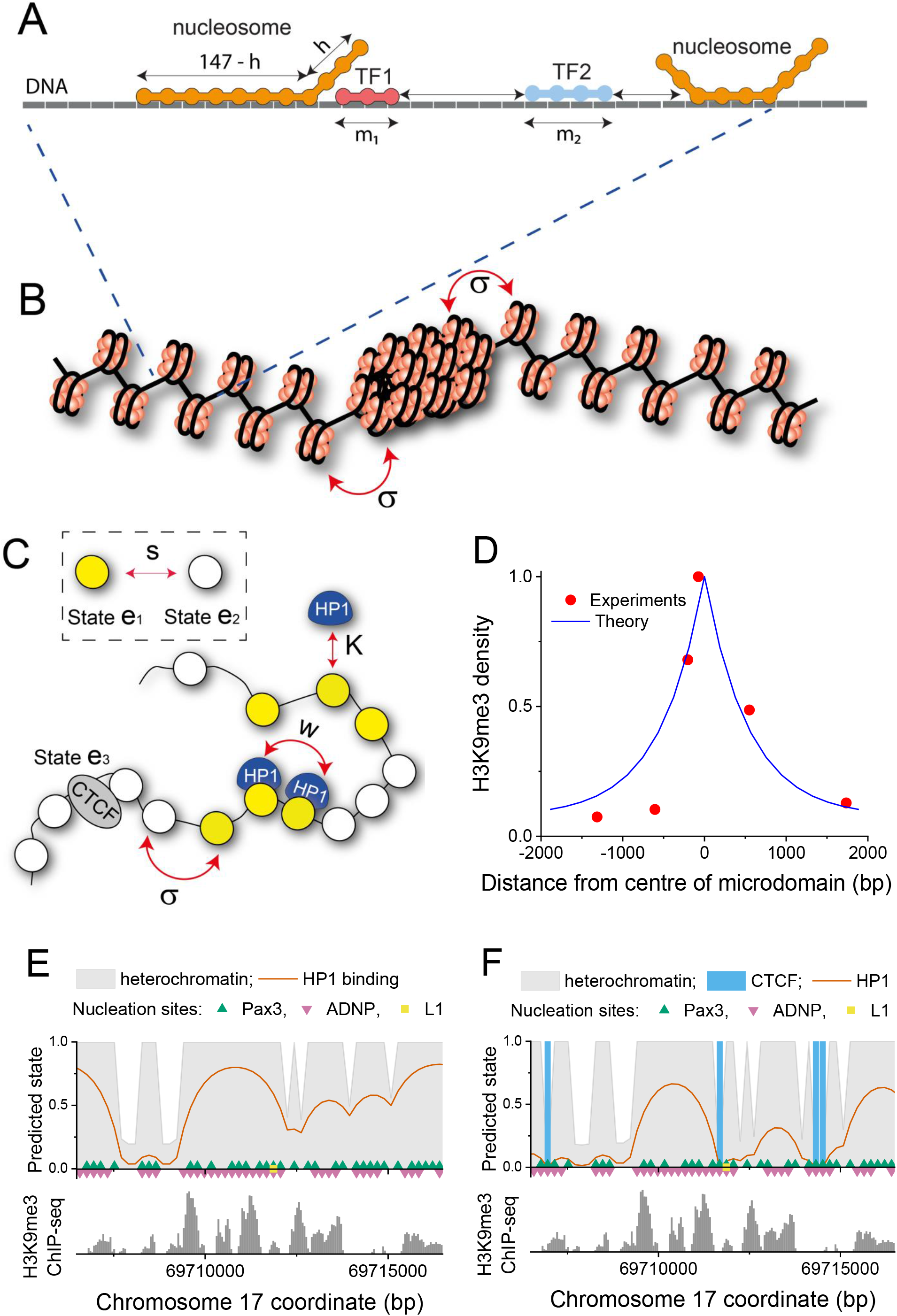
ChromHL framework for HND description. (**A**) A lattice model with single-base pair resolution for TF binding to DNA in the context of nucleosomes that can partially unwrap. (**B**) Schematic representation of the nucleosome array containing two types of packing. The boundary between different chromatin states is characterised by the statistical weight parameter σ. It could reflect that inter-nucleosome interactions in different packing states are different and thus create a boundary between them. (**C**) Description of chromatin in ChromHL. Chromatin is described by a lattice of nucleosome units. These lattice units can exist in different states. In the present study, three states e_1_, e_2_ and e_3_ exist that correspond to a H3K9me2/3 modified nucleosome, an unmodified nucleosome or a unit without a nucleosome but with CTCF bound. Lattice units can switch between different states with probability *s*(*e*_1_, *e*_2_). Chromatin proteins can shift this equilibrium by binding nucleosomes with different binding constants *K*(*e,g*) depending on the protein type *g* and the chromatin state *e* of a given lattice unit while interactions between proteins bound to neighbouring lattice units is described by the cooperativity parameter *w*. (**D**) H3K9me3 profile predicted by ChromHL in comparison to experimentally determined data in mouse embryonic fibroblasts for an ectopic HND^4^. (**E**) Model predictions (top panels) and experimental ChIP-Seq profiles of endogenous H3K9me3 (bottom panels) for an example genomic region in ESCs. This modelling takes into account HND nucleation at PAX3/9, ADNP and L1 sequence motifs, but does not consider CTCF binding. (**F**) Same as panel E but including CTCF binding as a factor that determines HND boundaries. It can be seen that including CTCF improves the predicted HND profile.

### An ectopically induced HND is confined by boundary interactions of nucleosomes

We first applied the ChromHL model to an artificial system of a single HND with a well-defined nucleation point and no sequence-defined boundaries. Such an ectopic HND was created in experiments of Hathaway et al. by tethering HP1α to the *Oct4* locus and inducing local H3K9me3 enrichment^4^. The experimentally determined H3K9me3 profiles decay to zero at distances of ~2,000 bp from the initiation site. We performed an optimisation with ChromHL to match this experimental H3K9me3 profile (**Fig. 2D, Fig. S4**). The heterochromatin spreading for this system is determined by three parameters: (i) The nucleosome binding activity of HP1 to H3K9me3-modified nucleosomes was estimated to be 10-fold stronger than that to unmodified H3K9 following previous publications^18^. (ii) The contact cooperativity value *w* for HP1-HP1 interaction has not been well defined in previous studies^18,43,44^. The best fit of the model to the H3K9me3 profile reported by Hathaway et al returned value of w = 4060. It is indicative of a significant positive binding cooperativity as compared to w = 1, which would represent independent binding of HP1 to adjacent nucleosomes. (iii) The parameter σ describes the “energetic boundary” between neighbouring chromatin packing types “e_1_” and “e_2_”. A value of σ = 1 would mean that this transition is not associated with energetic costs. However, for the ectopic HND our best fit required σ ~10^-5^. The magnitude of this σ-value is characteristic for highly cooperative transitions such as, for example, DNA melting^45^. The low value of σ ~10^-5^ means that the domains become intrinsically confined without additional DNA sequence-dependent contributions. This behaviour is different from endogenous HNDs considered below.

### ChromHL predicts experimental maps of endogenous HNDs in living cells

The nucleation sites of endogenous HNDs are determined by the genomic location of PAX3/9, ADNP and L1 sequence motifs derived above. When combined, they allowed a good match between computationally predicted and experimental HND profiles as shown for an exemplary region in ESCs (**Fig. 2E, F**). In addition, taking into account CTCF binding led to even better match of theory and experiment (**Fig. 2F**) as opposed to the model without CTCF (**Fig. 2E, Fig. S5**). Thus, endogenous HNDs depend to a larger degree on the DNA sequence than the ectopic HND (**Fig. 2D**). On the other hand, adding strong nucleosome-nucleosome interactions with σ ~10^-5^ as in the ectopic example leads in the case of endogenous HNDs to merging of the neighbouring nanodomains, while the fine structure of the H3K9me3 profile is lost (**Fig. S5, Fig. S7**). In the case of the endogenous SUV39H HNDs, a better fit was obtained with σ ~1. This means that the energy of nucleosome-nucleosome interactions at HND boundary does not exhibit any abrupt change, and CTCF binding is the main determinant of boundary formation. This model results in a larger number of smaller HNDs as the DNA sequence introduces many additional constrains to HND sizes (**Fig. S5, S7**). Thus, ChromHL allows us to separate different contributions of genetic and epigenetic interactions to the domain boundaries.

### Average endogenous nanodomain profiles have a typical extension of 0.7-2 kb

The characteristic aggregated H3K9me3 profiles of SUV39H-, GLP- ADNP- and ATRX-dependent HNDs in ESCs are shown in **Fig. 3**. These experimental profiles were obtained by averaging all individual regions with the corresponding heterochromatin subtypes centred at the summits of ChIP-seq peaks (H3K9me3 in the case of SUV39H, ADNP and ATRX, and H3K9me2 in the case of GLP). The resulting profiles resemble that of the ectopically induced H3K9me3 domain (**Fig. 2D**). Accordingly, the computational analysis of the aggregated data with the ChromHL model yielded a very good fit to the same model that was used for the ectopic HND: a central nucleation site and self-contained extension due to an unfavourable chromatin state transition as reflected by a low σ value (**Fig. 3, Fig. S4**). However, a closer inspection of the data reveals a significant variation of σ with a relatively high value of σ = 0.14 retrieved for SUV39H HNDs. Importantly, the information about molecular mechanisms that define the boundaries for individual regions is lost in the aggregated plots. Therefore, in the next part of this study we perform genome-wide analysis of individual domains.

**Fig. 3.**
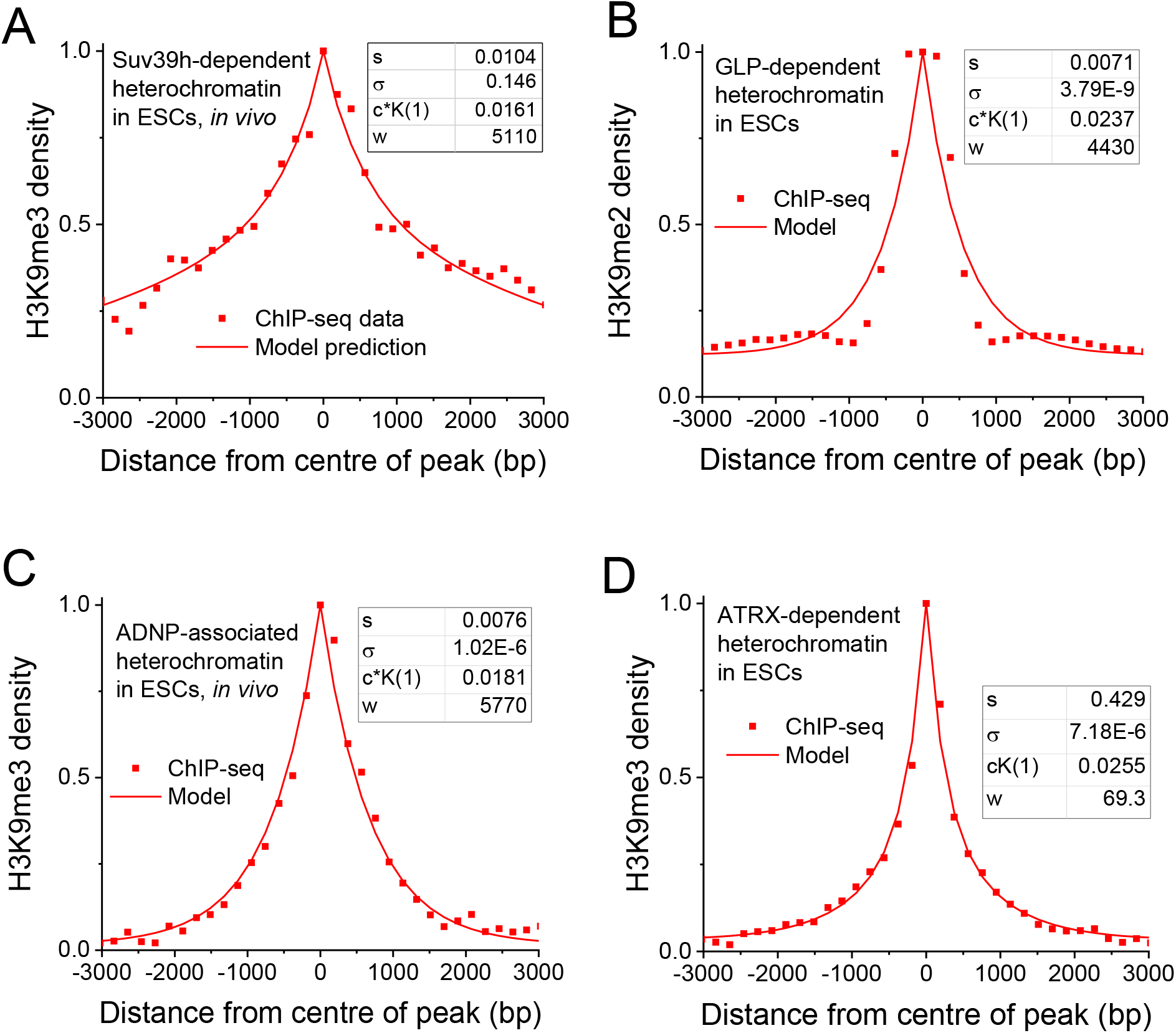
Shape of averaged H3K9me2/3 profiles for different HND types. The experimental data points were normalised to [0, 1]. s is the statistical weight for the nucleosome conversion to heterochromatin state, σ = σ(1,2) is the weight for interaction between nucleosomes in states 1 and 2, c·K(1) is the product of the local concentration of HP1 proteins in nucleosome clutch in state 1 and the binding constant of HP1 for this state. (**A**) SUV39H-dependent HNDs. (**B**) GLP-dependent HNDs. (**C**) ADNP-associated HNDs. (**D**) ATRX-dependent HNDs.

### DNA sequence is a major determinant of endogenous HNDs

Next, we investigated the effect of DNA sequence on heterochromatin initiation and localisation. A genome-wide analysis was conducted for the four different HND types with the nucleation sequence motifs derived above. In our analysis we considered both the effect of CTCF and cooperative HP1 binding to neighbouring nucleosomes with a 10-fold increase of the binding constant at H3K9me2/3-modified nucleosomes (**Fig. 4, Table 2**). The comparison of predicted and experimentally determined distributions of SUV39H-dependent HND sizes showed a significant improvement if CTCF binding was included and yielded a fit value of σ = 1 (**Fig. 4A, B, Supplemental Fig. S6, S8A, B**). Interestingly, the model derived for SUV39H-HDNs was also well suited to describe the GLP-HNDs marked by H3K9me2 (**Fig. 4B, Fig. S8B**). For ADNP-HNDs we used the PWM derived from ADNP ChIP-seq to define ADNP binding sites for domain nucleation (**Fig. 4C**). Again, including CTCF binding improved the model with σ = 1. For ATRX-HNDs, L1 repeats were used as nucleation sites. The resulting model describes the experimental data well (**Fig. 4D, Fig. S8D**). Notably, and in contrast to the three other heterochromatin types, the effect of CTCF was negligible. Furthermore, the best fit value of the boundary weight yielded σ = 0.01, corresponding to a free-energy change ≈ 4.6 *kT*. This energy is comparable to typical nucleosome-nucleosome interactions^18^ and very different from the value of σ = 1 (energy change ≈ 0 *kT*) obtained as best fit for the other heterochromatin types. Thus, we conclude that boundaries of ATRX-HNDs are determined mostly by unfavourable transitions to the flanking chromatin states. In contrast, the SUV39H-, GLP- and ADNP-HNDs were described best with the same model that included sequence specific binding of TFs (PAX3/9, ADNP) as a nucleation site, CTCF binding sites as boundary elements and a value of σ = 1 indicative of no energy penalty to the flanking chromatin states.

**Fig. 4.**
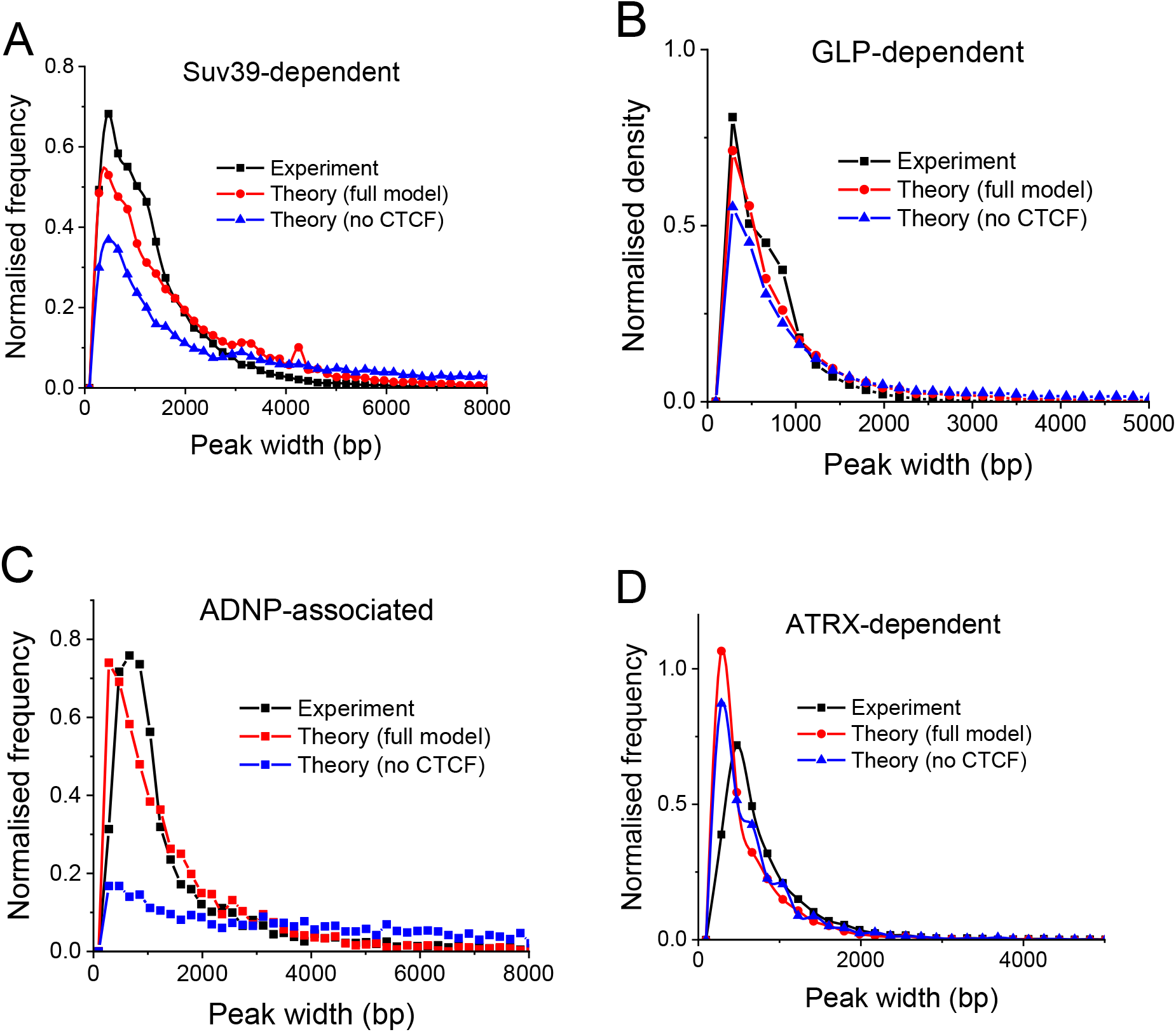
ChromHL predictions compared with the experimental distribution of HND sizes. Panels on the left show the histograms of the experimental nanodomain sizes given by H3K9-methylated ChIP-seq peaks (black) against the theoretically predicted ones using the full model (red) and the simplified model that does not take into account CTCF (blue). Panels on the right show the correlation between the nanodomain sizes and the number of heterochromatin-initiating motifs per nanodomain. Parameters of the best fit obtained with ChromHL are given in **Table 2**. (**A**) SUV39H-dependent HNDs. (**B**) GLP-dependent HNDs. (**C**) ADNP-associated HNDs. (**D**) ATRX-dependent HNDs computed with the L1 repeats-based model and *σ* = 0.01.

**Table 2.** Parameters of ChromHL models for the four types of HNDs

|  | **SUV39H<sup>a</sup>** | **GLP<sup>a</sup>** | **ADNP<sup>a</sup>** | **ATRX<sup>a</sup>** |
| --- | --- | --- | --- | --- |
| **Nucleation factor** | PAX3/9 | PAX3/9 | ADNP | LINE1 binder |
| **HND initiation log threshold** | -4.5 | -4.0 | -9.5 | No threshold |
| **s** | 0.2 | 0.2 | 0.2 | 0.2 |
| **σ** | 1.00 | 1.00 | 1.00 | 0.01 |
| **HP1 cooperativity (w)** | 100 | 100 | 100 | 100 |

<sup>a</sup> Protein factor upon which the HNDs are dependent in ESCs.
<sup>b</sup> Log-affinity threshold for the binding of the nucleation factor.

### HNDs differ in their nucleosome packing patterns

We further dissected the differences between SUV39H-, GLP-, ADNP- and ATRX-HNDs by assessing the nucleosome repeat length (NRL) inside these regions. We calculated average NRLs using MNase-seq data based on cutting DNA between nucleosomes^46^ (**Fig. 5A**) as well as the dyad-to-dyad frequency distribution using chemical mapping data based on cutting DNA at the nucleosome dyads^47^ (**Fig. 5B**). MNase-seq derived NRLs for SUV39H-, GLP-, and ATRX-dependent nanodomains were similar to the genome-wide NRL of 189±1 bp. In addition, heterochromatin states defined previously based on H3K27me3 enrichment in ESCs using ChromHMM^48^ had a similar NRL value (**Fig. S11A**). In contrast, ADNP-associated HNDs were characterised by a smaller NRL of 175±1 bp. When considering the distribution of nucleosome dyad-to-dyad distances obtained from chemical cleavage at nucleosome dyads^47^ (**Fig. 5B**), SUV39H- and GLP-dependent heterochromatins again showed the same distribution as genome-average. In contrast, ATRX- and ADNP-HNDs clearly displayed a different distribution of dyad-to-dyad distances (**Fig. 5B**). ADNP-HNDs had significantly smaller dyad-to-dyad distances while the distribution of ATRX-HNDs was shifted to larger values in comparison to genome average. Interestingly, in SUV39H- and GLP-HNDs, the distribution of dyad-to-dyad distances resembled that of H3K27me-enriched heterochromatin (**Fig S11A**). This is consistent with the fact that ~75% of SUV39H- and ~90% of GLP-dependent HNDs reside within H3K27me-enriched chromatin states. In contrast, only about 5% of ATRX-dependent HNDs were located within H3K27me3-enriched states^48^ (**Fig. S11B**). We have studied the effect of different sizes of the ChromHL lattice unit on our predictions of the distribution of HND sizes, considered effective NRLs ranging from 161-199 bp, but did not observe significant effects of the NRL change on the nanodomain sizes *per se* (**Fig. S10**). This suggests that the differences in nucleosome packing found above may affect HND formation indirectly, e.g. by modulating the value of σ.

**Fig. 5.**
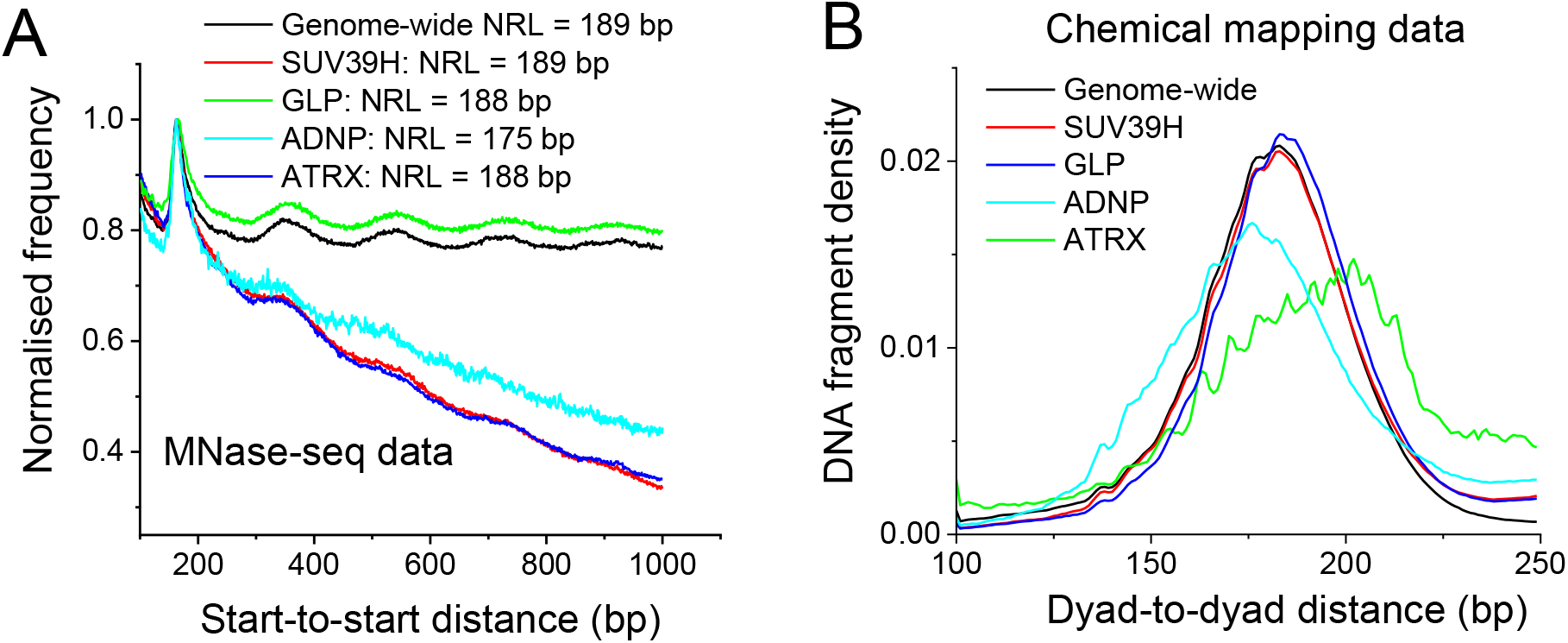
Nucleosome packing characteristics in different types of heterochromatin. (**A**) Nucleosome start-to-start genome-wide distance distribution (black line) based on MNase-seq^60^ in SUV39H- (red), GLP- (green), ADNP- (light blue) and ATRX-HNDs (dark blue). (**B**) Nucleosome dyad-to-dyad distance distribution for nearest-neighbour nucleosomes based on chemical mapping^47^.

### HND redistribution during cell transition can be regulated by protein binding activity

H3K9me2/3 marks constitutive heterochromatin loci as well as cell type specific regions that change during differentiation^49,50^. Furthermore, aberrant gain or loss of H3K9me3 is a feature of many cancers^51,52^. In our framework, the location and extension of HNDs is regulated by binding of PAX3/9, ADNP and CTCF. In general, the binding activity of these and other TFs can be regulated via their expression levels, subcellular localization and/or posttranslational TF modifications^35^. In addition, the activity of H3K9-modifying enzymes (SUV39H1/2, GLP, SETDB1) or H3K9me2/3-binding proteins like HP1 could determine cell type specific HND patterns. Significant changes in the binding properties and genomic localization of HP1 molecules occur during cell differentiation^53,54^. Accordingly, we explored the effect of the change of HP1 concentration on the structure for SUV39H-dependent H3K9me3 HNDs (**Fig. 6A**). Our model predicts that decreasing the concentration of free HP1 molecules reduces HND size. Consistent with this prediction, the experimental distributions of SUV39H-dependent HNDs in ESCs vs neural progenitor cells (NPCs) generated by *in vitro* differentiation shows a significant decrease of average HND size (**Fig. 6B**). Similar effect takes place for all four types of HNDs upon differentiation of ESCs to NPCs (**Fig. S12**). HND shrinking/expansion in dependence of HP1 activity can be further modulated by differential binding of HND-initiating TFs and CTCF as was shown above (**Fig. 2E, Fig. 2F**). Thus, the formation of HNDs is partially hard-wired in the DNA sequence but their cell-type specific patterns are dependent on the activities of additional factors (**Fig. 6C**).

**Fig. 6.**
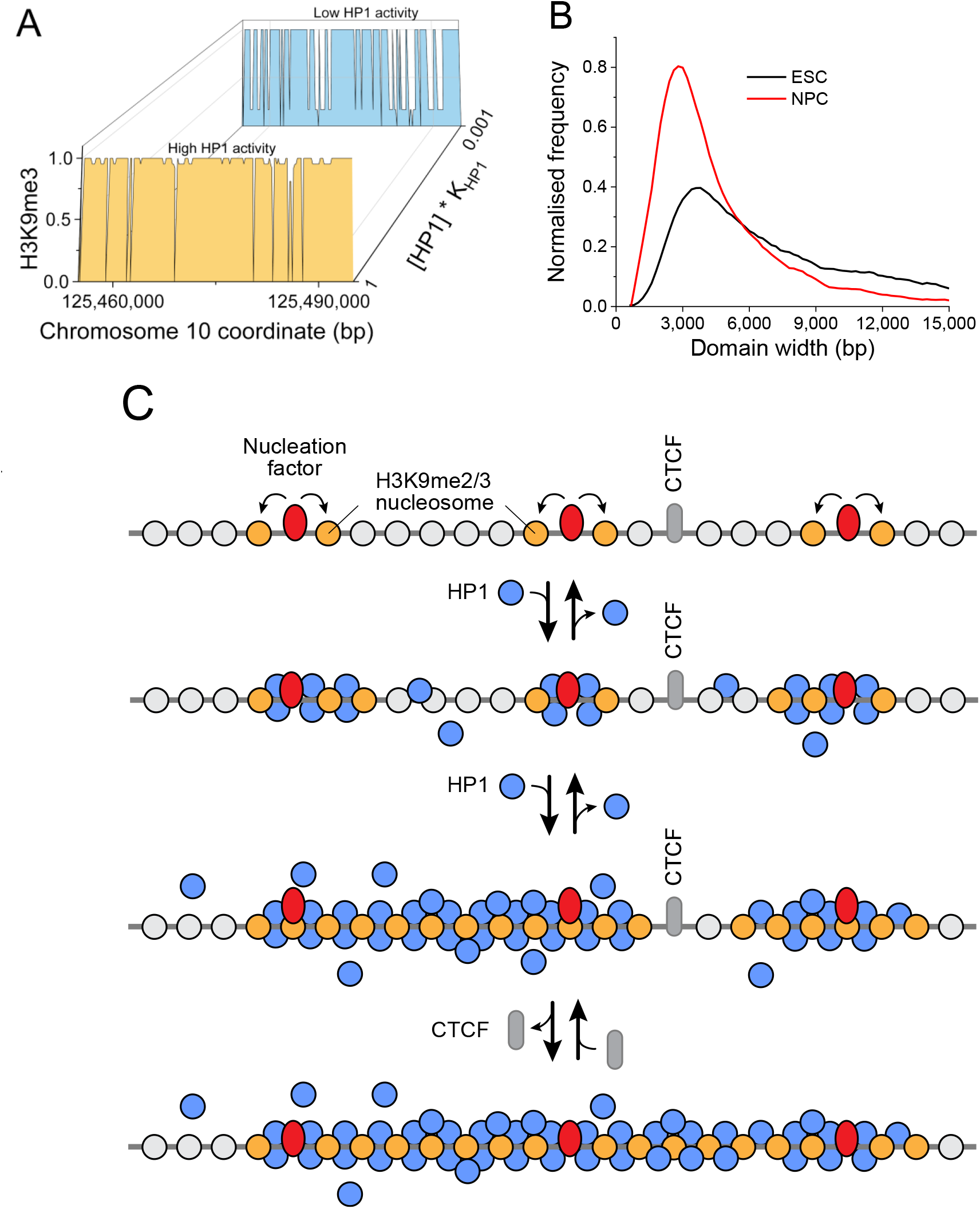
Dynamics of heterochromatin nanodomains during cell transitions. (**A**) Model-predicted heterochromatin state probability for the region centred on chr10:125454974-125464974, for large ([HP1] × K_HP1_ = 1) and low concentration of free HP1 molecules ([HP1] × K_HP1_ = 0.001). The model predicts that increasing concentration of free HP1 molecules leads to formation of larger heterochromatin domains. (**B**) Experimental distributions of H3K9me3 domains in ESCs (black) and NPCs differentiated from them (red). (**C**) A schematic model of heterochromatin domain dynamics. Heterochromatin nucleation sites are determined by the DNA sequence and concentration of molecules binding these sequences, such as Pax3/9 and ADNP. Decreasing the HP1 activity (the local concentration of free HP1 molecules) may lead to global narrowing heterochromatin domains, as observed during ESC differentiation to NPCs. In addition, changes of CTCF binding leads to changes of heterochromatin domains boundaries.

## Discussion

We conducted a systematic comparison of four types of HNDs in ESCs and quantitatively described them with ChromHL, the first framework allowing sequence-specific prediction of HND maps. The SUV39H- and GLP-HNDs were well-described by a common model based on PAX3/PAX9-nucleation sites (**Fig. 4A**). Binding of PAX3/9 has been previously suggested to target SUV39H-dependent H3K9me3 domains^29,30^. Our analysis supports this conclusion for the corresponding HNDs as we find a high correlation between the number of PAX3/9 motifs per domain and domain size (**Fig. 1C, Fig. 1F, Table 1, 2**). Interestingly, the same model worked for GLP-dependent HNDs that carried the H3K9me2 mark. The ADNP-associated HNDs could also be well described with a ChromHL model using the ADNP motif computed here (**Fig. 4C, Table 1, 2**). It is noted that the ChromHL analysis is based on the presence of sequence motifs and thus it is possible that additional TFs are involved as nucleating factors that recognise these sequences. This consideration is particularly relevant for the case of ATRX-dependent HNDs. Our analysis identified L1 repeat family as the best sequence feature responsible for HND nucleation (**Fig. 1L**). However, the correlation of the sizes of ATRX-HNDs with the number of L1 motifs per HND was only 67% and the use of this sequence feature to model ATRX nanodomain formation with ChromHL resulted in a less perfect fit in comparison to the other three HNDs (Fig. 4D). Thus, additional protein factors and nucleation mechanisms are likely at play for ATRX-dependent HNDs. It’s also worth noting that ATRX knockout reduces H3K9me3 on IAPs, but H3K9me3 is not lost entirely, so these regions would not qualify as ATRX-dependent HNDs in the above analysis, but still ATRX may have effects on these regions.

Our ChromHL analysis showed that CTCF binding motifs at boundaries represent a major defining feature for the extension of SUV39H-, GLP- and ADNP-HNDs. This contribution of CTCF is consistent with our recent report where CTCF sites acted as bifurcation points for differential DNA methylation spreading upon TET1/2 knockout^26^. CTCF effect on the spreading of H3K9 methylation is a novel aspect arising from the current analysis. CTCF is known to be involved in formation of topologically associated domains and loops, with the size of CTCF-demarcated loops/and in the range from 10 kb to 1 Mb^55^. In contrast, the nanodomains studied here are typically 0.7-2 kb in size and involve weak CTCF binding sites that are frequently not called with the typical peak detection thresholds used in the analysis of CTCF ChIP-seq data that retrieve ~60,000 relatively strong binding sites. However, we propose here that weak CTCF sites are functionally important and define the H3K9me2/3 nanodomain structure in the genome by transient binding of CTCF, possibly in conjunction with other proteins. In our recent work, we reported that such CTCF motifs are enriched in DNA sequence repeats at sites of reduced nucleosome density^56^. Thus, the effect of these motifs may also involve loss of nucleosomes, which could affect the interactions between neighbouring nucleosomes at the boundary.

ChromHL modelling allowed us to uncouple DNA sequence determinants from thermodynamic constraints that limit the sizes of HNDs. The parameter σ defines the energetic cost of formation of a new boundary between chromatin states in analogy to the cooperativity constant used in statistical mechanical models that describe melting of the DNA double helix^45^. The associated boundary energies can, for example, arise from (un)favourable nucleosome stacking interactions between nucleosomes^57^. In the case of ectopic HNDs established in the experiments of Hathaway et al^4^, the best fit of our model returned small σ values suggesting unfavourable nucleosome interaction energies at the domain boundaries. In contrast, in the case of endogenous HNDs, the best fit for the SUV39H-, GLP- and ADNP-nanodomains was obtained with a boundary formation weight of σ = 1, indicating a lack of structural transitions from the H3K9me2/3 states. This finding suggests that the SUV39H-, GLP- and ADNP-dependent nanodomain size is mostly DNA sequence-determined by the respective nucleating TFs and CTCF. In contrast, ATRX-dependent H3K9me3 nanodomains did not fit to this model. A value of σ = 0.01 was retrieved that corresponds to a significant energetic cost of nanodomain boundary formation. In line with this observation, nucleosome occupancy and distribution in ATRX-dependent HNDs were significantly different from other types of HNDs (**Fig. 1D, 5B**). Thus, thermodynamics of nucleosome packing plays a more important role in limiting the ATRX-dependent nanodomain size.

Our model revealed an important role of the DNA sequence as a determinant of HND formation. This raises the question, whether these domains are epigenetically regulated or represent mostly constitutive heterochromatin. Since ChromHL explicitly includes chromatin ligand binding, one straightforward mechanism for cell type specific HND formation would be to epigenetically regulate the activity/concentration of the nucleation factors, CTCF or histone methylases that set the H3K9me2/3 mark. In addition, we showed here how the change of the HP1 concentration and corresponding nucleosome occupancy induces the shrinking/merging of nanodomains, which could drive cell-type specific differences (**Figs. 6B, 6A, S12**). Thus, by modulating the local concentrations of proteins that initiate (e.g., PAX3/9, ADNP), stop (e.g., CTCF) or promote the spreading of chromatin nanodomains (e.g., HP1), the cell can regulate the epigenetic states that are otherwise pre-determined by the DNA sequence (**Fig. 6C**). In addition, ADNP and CTCF can compete for binding sites^58^, which could contribute to a modulation of ADNP domain size extension in dependence of the ADNP/CTCF binding activity ratio. CTCF (and other TFs) also bind in competition with nucleosomes that can adopt different positions as reflected in the NRL analysis conducted here. Accordingly, cell-type specific binding of CTCF to certain sites is driven by a complex interplay of nucleosome binding, DNA (de)methylation and other factors^26,59^.

In summary, the ChromHL modelling approach developed here identified key parameters for the description of HNDs that are abundant in the genome. The analysis has been conducted for mouse embryonic stem cells but can be applied to other cell types and chromatin nanodomain types as well. We anticipate that it will be valuable to distinguish sequence- and non-sequence-dependent epigenetic effects for establishing cell type specific gene expression programs.

## Methods

### Cell lines and cell culture work

Wild type murine embryonic stem cells (ESCs) wt26 and *Atrx* knock out cell lines (KO1-40 and KO1-45) were described previously^36^. Cells were cultured on 0.2% v/v gelatine (in PBS) in high glucose DMEM (Gibco 31053-028) supplemented with 1 mM sodium pyruvate and 4 mM L-glutamine (PAA M11-006), 15% v/v FCS (Sigma F7524, lot: 091M3398), 1% v/v penicillin-streptomycin (PAN Biotech P06-07100), 100 μM β-mercaptoethanol (Sigma 63689), 1% v/v non-essential amino acids and 0.41% v/v LIF (self-made; supernatant from LIF-producing cells; batch: 7/26/14).

### ChIP-seq to map ATRX HNDs

ChIP-seq experiments were conducted essentially as described before (Teif et al. 2012). To shear the chromatin, cells were digested with MNase for 15 minutes in a buffer containing 25 mM KCl, 4 mM MgCl_2_, 1 mM CaCl_2_, 50 mM Tris/HCl pH7.4 and 1x protease inhibitor from Cell Signalling and sonicated with a Covaris S2 sonicator (parameters: 900 s, burst 200, cycle 20%, intensity 8) in sonication buffer (10 mM Tris pH 8.0, 200 mM NaCl, 1 mM EDTA, 0.5% N-lauroylsarcosine, 0.1% Na-deoxycholate). For the pre-clearance, 4 μg normal rabbit IgG (R&D Systems, AB-105-C, lot: ER1212071) and ChIP-grade protein G magnetic beads (Cell Signalling 9006S, 25 μl/sample) were used. A 1/20 fraction of the supernatant was used as input sample and the remaining material was split for three IP reactions with an anti-H3K9me3 antibody (Abcam, ab8898, lot: GR148830-2). An amount of 4 μg antibody was added to each IP sample, incubated at 4°C for 2 h, protein G magnetic beads were added and then the mixture was incubated at 4°C overnight. After elution of the IP samples, cross-linking was reversed, RNase A and Proteinase K digestion were added, and DNA was precipitated. Experiments were conducted for two replicates of the wild-type cell line (wt26) and one replicate of each *Atrx* ko cell line (KO1-40 and KO1-45). Libraries for ChIP-seq were prepared with the NEBNext Ultra DNA Library Prep Kit for Illumina (New England Biolabs, NEB #E7370) according to the manufacturer’s instructions. In the size selection step, insert fragments of 150 bp corresponding to 270 bp total library size (including insert and adaptors) were selected. A total of 13 PCR amplification cycles were carried out and the library size and quality were checked by gel electrophoresis. All samples were sequenced on an Illumina HiSeq 2000 platform at the DKFZ Sequencing Core Facility.

### Peak calling of ChIP-seq data

For a given type of HND, the differential H3K9me3 or H3K9me2 peaks were called with MACS2 (Zhang et al. 2008) of WT and the KO datasets of *Suv39h1/h2, Glp*, and *ATRX,* respectively, against the common input, using the parameter broad-cutoff 0.1. Peaks present in WT but not in the KO cells were retained as peaks that were dependent on a given factor. In the case of ADNP-associated HNDs a H3K9me3 dataset in *Adnp^-/-^* cells was not reported^32^. Therefore, we have defined ADNP-associated HNDs as the intersection of ADNP-bound ChIP-seq peaks with all H3K9me3 peaks in wild type ESCs from these experiments (*n* = **4,673**). Manipulations with BED files were performed using bedTools^61^. For H3K9me3 in NPCs, we used datasets GSE61874^54^ and GSE57092^30^ with peak calling performed by MACS for Figure S12 as well as EPIC^62^ for Figure 6B. We then intersected H3K9me3 peaks in ESCs and NPCs and retained in the analysis only those peaks which overlapped between these two conditions.

### Analysis of chromatin features

Our chromatin annotation used the 15 chromatin states assigned to ESC genomic regions previously^48^ using the package ChromHMM^63^. The nucleosome repeat length (NRL) was determined based on the previously published MNase-seq dataset^60^ using NucTools as described previously^46,59^. The dyad-dyad differences were computed using the chemical mapping dataset^47^. The Kolmogorov-Smirnov test p-values for the histograms of the dyad-dyad distance distributions were calculated using OriginPro software (OriginLab). Nucleosome occupancy, CTCF binding occupancy, HP1 binding, mappability, CpG methylation, H3K9me3, H3K9me2, H3K4me1, H3K27me3 were computed on regions of 40,000 bp centred on the nanodomain centre using NucTools^46^ and averaged. These were further smoothed using a 2,000 bp Savitsky-Golay filter of order 2. Enrichments were computed with the BedTools as the ratio of observed number of intersections between a given datasets and a given genomic feature to the number expected by chance for the same number of randomly shuffled regions.

### Identification of nucleation sites

In the case of SUV39H- and GLP-dependent heterochromatin we scanned the DNA sequences using RSAT^64^ with the PAX3 and PAX9 position weight matrix (PWM) obtained from TRANSFAC^65^ to obtain the locations of the motif within each peak. In the case of ATRX-dependent heterochromatin we obtained the list of specific L1 peaks (L1Md_F2, L1Md_T, L1Md_A and L1Md_F) using the RepeatMasker tool from the UCSC Genome Browser^66^. In the case of ADNP-associated HNDs we derived the PWM for sequencespecific ADNP binding using MEME^67^ based on **100-bp summits of 600 top** ADNP-bound ChIP-seq peaks from Ostapcuk et al^32^. This PWM was then used for DNA sequence scanning with RSAT to determine ADNP motif locations inside H3K9me3 ChIP-seq peaks intersecting with ADNP-bound ChIP-seq peaks. Telomeric repeats were defined as single telomeric repeat motif reported^68^ using RSAT^64^. G-quadruplex repeats were defined as regex search on the sequences for each ATRX dependent peak with fastaRegexFinder by Dario Beraldi to identify the DNA sequence motif (G_3_N_1-7_)_3_G_3_ (https://github.com/dariober/bioinformatics-cafe/tree/master/fastaRegexFinder).

### ChromHL modelling

In the calculations, we distinguished three “epigenetic” states (Fig. 1): heterochromatin (*e* = 1), euchromatin (*e* = 2) and the absence of nucleosome due to CTCF binding (*e* = 3). Outside of any CTCF binding or heterochromatin initiation site, we set *s*(*i*, 1) = *s*; *s*(*i*, 2) = 1; *s*(*i*, 3) = 0. The initiation of heterochromatin was specified in the model by setting *s*(*i*, 1) = 1, *s*(*i*, 2) = 0, *s*(*i*, 3) = 0, for the lattice unit *i* where initiation occurs. For the HP1 *in vitro* recruitment experiment, this was only at the centre point of the lattice. For the weights of nucleosome-nucleosome contacts, σ(e_1_, e_2_) = σ_e1e2_, we focus on the single most important boundary parameter σ12 = σ21 = σ and set all other contact weights equal to 1. We also set all cooperativity parameters equal to 1, except for the contact HP1-HP1 cooperativity (**Fig. S9**). The latter parameter was fitted in the case of the Hathaway et al experiments, or fixed in the case of *in vivo* heterochromatin formation to *w*(1,1,0) = 100 or *w*(1,1,0) = 1, as explained in the corresponding figures. For TF-based heterochromatin initiation models the TRAP affinity score^69^ was computed using the corresponding PWM based on the DNA sequence. This affinity was then geometrically averaged over a 501-bp window and subjected to a threshold for including a lattice unit as a nucleation site (**Supplemental Fig. S13, S14**). For SUV39H-dependent HNDs, we computed receiver-operator (ROC) curves using smoothed affinity scores on the peaks and matched non-peaks for different smoothing windows sizes (**Supplemental Fig. S2B, S2D**). The maximum area under the ROC curve (AUC) was for 1001-bp windows. However, the AUC for this window size is not significantly different to that for 501bp (**Supplemental Fig. S2C, S2E**) and using that window size smooths the affinity profiles too much, worsening the theory-experiment fit (data not shown). Thus, we used a 501 bp geometric mean centred smoothing window for determining heterochromatin initiation sites for these sets of peaks, assuming PAX3/9 binding is the initiation factor. For the ATRX-dependent heterochromatin, L1 sub-repeats L1Md_F2, L1Md_T, L1Md_A and L1Md_F that were found to be overrepresented in the ATRX-dependent H3K9me3 peaks were downloaded from the RepeatMasker track on the UCSC Genome Browser (**Supplemental Table S1**). A lattice unit containing the centre of a repeat was considered an initiation site. The length of the lattice in units was calculated by placing a lattice unit exactly in the centre of the region and adding units either side of this position symmetrically until all the sequence was covered.

## Data and software availability

Data from the ATRX knockout experiments is deposited in the GEO database under accession number GSE158744. The ChromHL software and associated codes are available at https://github.com/TeifLab/ChromHL.

## Acknowledgements

We thank Thomas Manke, Victor Zhurkin and Thomas Höfer for fruitful discussions. This work was supported by the Wellcome Trust grant 200733/Z/16/Z to VBT and by grant RI1283/16-1 to KR in the DFG priority program 2191.

