## Supplemental Materials for "DNA sequence-dependent formation of heterochromatin nanodomains"

### Theory

Our proposed theoretical model is motivated by classical Ising-type models [1] as was traditionally employed in DNA-binding and DNA-melting studies [2,3], considering chromatin as a quasi-1D lattice of units. Specifically, we base our model on the general transfer matrix formalism [4-7], extended to take into account different scales from nucleotide- to nucleosome- to chromatin domain-resolution, aka hierarchy. This hierarchy of scales means that that we need define the lattice separately at the level of DNA base-pairs as lattice units (when sequence-specific TF binding is considered) and at the level of nucleosomes as lattice units (when chromatin state transitions are considered).

At the nucleosome-lattice level we assume that the interaction between the neighboring nucleosomes depends on the characteristic NRLs, as well as other structural effects which are all cast in the phenomenological definition of the "chromatin state" (Figure 1). A nucleosome can belong to any chromatin state, characterized by a certain self-energy depending on the given state, a nucleosome-nucleosome interaction energy depending of the states of the two nucleosomes, and a protein binding energy depending on the protein type and the nucleosome state. Proteins bound to neighboring nucleosomes can interact with each other. Previously we have constructed a mechanistic model of heterochromatin protein 1 (HP1) binding to the nucleosome array [8] based on *in vitro* binding experiments [9]. Here we add to the HP1 model additional players such as CTCF, Pax3, Pax9 and ADNP capable to initiate heterochromatin nanodomains. We parameterize this model based on the published experimental datasets for mouse embryonic stem cells (ESCs) with respect to targeted recruitment of HP1 [10], Suv39-dependent H3K9me3-marked heterochromatin [11], GLP-dependent H3K9me3-marked heterochromatin [12], our original experiments reported here with respect to ATRX-dependent H3K9me3 heterochromatin studied in ESCs with knockout of ATRX enzyme, as well as a recent work reporting ADNP-dependent H3K9me3 heterochromatin [13].

Let us base our solution of the lattice model on the general transfer matrix formalism [4,5,7,8]. In this method each elementary lattice unit can be in a number of states and it can be affected only by a limited number of neighboring lattice units (e.g. just the next neighbor unit in the limiting case of contact cooperativity, or up to V next neighbor units in the more general case accounted for in our previous publications [7,14-16]). All states need to be enumerated and the corresponding statistical weights need to be assigned for all combinations of allowed states of a lattice unite number *i* given the state of the next unit *i*+1. The matrix that stores these weights is called the transfer matrix. In particular, in our previous publication a transfer matrix called *MatrixUnwrap* has been already constructed at the single-base pair level for a model that takes into account cooperative competitive TF binding to the DNA inside and outside the nucleosome and longer than nearest-neighbor interactions [7]. Here we increased the complexity of the model by introducing additional arbitrary states of the lattice (e.g. belonging to a nucleosome in a "heterochromatin" or "euchromatin" state, or belonging to a nucleosome-free "insulator region").

In order to explain the general matrix solution of this problem, let us first demonstrate this concept for a simplified situation. Let us consider non-sequence-specific protein binding to DNA that is characterized by contact cooperativity. In this case, each lattice unit can be in two states: bound or unbound, and the corresponding transfer matrix *A* for is 2x2:

$$A = \begin{pmatrix} wKc & Kc \\ 1 & 1 \end{pmatrix},$$

where each raw corresponds to the state of a given lattice unit *i*, and each column to the state of the next lattice unit *i*+1, *K* is the binding constant, *c* is the free protein concentration,

$w$ is the contact cooperativity between DNA-bound proteins. In this case we assigned 1 as the weight of a lattice unit that is not bound by protein. The weight of a lattice unit bound by the protein is $Kc$. The weight of a lattice unit bound by a protein interacting with the next lattice unit that is also bound by a protein is given as a product $w*K*c$, where $w$ is the contact cooperativity constant.

Let us also consider another transfer matrix $B$ corresponding to a simplified two-state nucleosome lattice. In this case each lattice unit can be in two states: say, heterochromatin, and euchromatin. Let us characterize each of these states by a weight $s_1$ and $s_2$ correspondingly. Let the interaction between two nucleosomes is characterized by a weight $\sigma$ that depends on the state of each of these two nucleosomes. In this case the transfer matrix $B$ will have a simple form:

$$B = \begin{pmatrix} s_1\sigma_{11} & s_1\sigma_{12} \\ s_2\sigma_{21} & s_2\sigma_{22} \end{pmatrix}$$

Now if we combine the two effects (protein binding reflected in matrix $A$ and nucleosome $s_1/s_2$ transition reflected in matrix $B$), each lattice unit will have four possible states (protein-bound $s_1$; unbound $s_1$, protein-bound $s_2$, unbound $s_2$), and the corresponding transfer matrix will $C$ will be 4x4 as summarized below:

$$C = \begin{pmatrix} w_{11}K_1c_1s_1\sigma_{11} & K_1c_1s_1\sigma_{11} & w_{12}K_1c_1s_1\sigma_{12} & K_1c_1s_1\sigma_{12} \\ s_1\sigma_{11} & s_1\sigma_{11} & s_1\sigma_{12} & s_1\sigma_{12} \\ w_{12}K_2c_2s_2\sigma_{21} & K_2c_2s_2\sigma_{21} & w_{22}K_2c_2s_2\sigma_{22} & K_2c_2s_2\sigma_{22} \\ s_2\sigma_{21} & s_2\sigma_{21} & s_2\sigma_{22} & s_2\sigma_{22} \end{pmatrix}$$

In principle, we can distinguish free protein concentration $c$ inside chromatin regions in state 1 and 2, denoted correspondingly $c_1$ and $c_2$. Different chromatin states can have different free concentrations of our protein of interest e.g. due to liquid-liquid phase separation in chromatin [17-19].

A system characterised by the simple transfer matrix above can be solved even analytically, deriving exact expressions for the average size of the heterochromatin nanodomain, average number of nanodomain boundaries, etc. However, in reality DNA-protein binding is sequence specific and depends on the chromatin states. Thus, the parameters $s$ and $K$ are the functions of the location of the lattice unit along the genomic coordinate ($n$), which precludes analytical solutions. To construct a DNA sequence-informed transfer matrix, we base our methodology on a more advanced transfer matrix *MatrixUnwrap* [7] instead of matrix $A$ shown above. The *MatrixUnwrap* model is defined for protein-DNA binding that takes into account binding of multiple types of proteins $g$ ($g$ = 1…$f$) at concentrations $c(g)$, which bind lattice units in different states $e_n$ with binding constants $K(n, g, e_n)$, and interact with each other with distance-dependent potential $w(g_1, g_2, l)$ ($l < V_g$). Furthermore, this transfer matrix takes into account that nucleosome can partially unwrap from the DNA. The calculation of the *MatrixUnwrap* elements has been described previously [7]. The total number of states in the *MatrixUnwrap* model is given by the following expression:

$$\sum_{g=1}^{f}\left(m_g + V_g\right) + 2 + \max(V_g) + 1$$

Here we introduce $e_{max}$ additional "epigenetic" states $e_n$ for each lattice unit ($e_n=[1,e_{max}]$). Correspondingly, each individual nucleosome is characterized by the weight $s(e_n) = \exp(\Delta G(e_n)/RT)$, where $\Delta G(e_n)$ is the energy of the nucleosome in state $e_n$, $R$ is the universal gas constant, $T$ is the temperature. Similarly, the contact between two neighboring nucleosomes in states $e_1$ and $e_2$ is assigned a weight $\sigma(e_{n,} e_2) = \exp(\Delta G(e_1, e_2)/RT)$, where $\Delta G(e_1, e_2)$ is the energy of nucleosome-nucleosome interaction. This results in the increase of the total number of states of the model as follows:

$$e_{\max} \cdot \left[ \sum_{g=1}^{f} (m_g + V_g) + 2 + \max(V_g) + 1 \right]$$

A detailed description of the assignment of all transfer matrix states is available in the open-source MATLAB code available at https://github.com/TeifLab/ChromHL.

Once sequence-dependent transfer matrices $Q_n$ are constructed for each lattice unit, we can calculate the partition function $Z$ of the lattice of length $N$ by multiplying all transfer matrices $Q_n$ sequentially:

$$Z = (1\ 1 \ldots 1) \times \prod_{n=1}^{N} Q_n \times \begin{pmatrix} 1 \\ 1 \\ \ldots \\ 1 \end{pmatrix}$$

The probability that a given lattice unit $n$ is in state $x$ can be calculated as

$$P_n(X) = \frac{s_{xn}}{NZ} \times \frac{\partial Z}{\partial s_{xn}},$$

where $s_{xn}$ is the statistical weight for a state x for a given lattice unit. In particular, we can calculate the binding maps for any protein of interest, as well as the maps of “epigenetic” states for any DNA sequence and concentrations of regulatory proteins such as HP1.

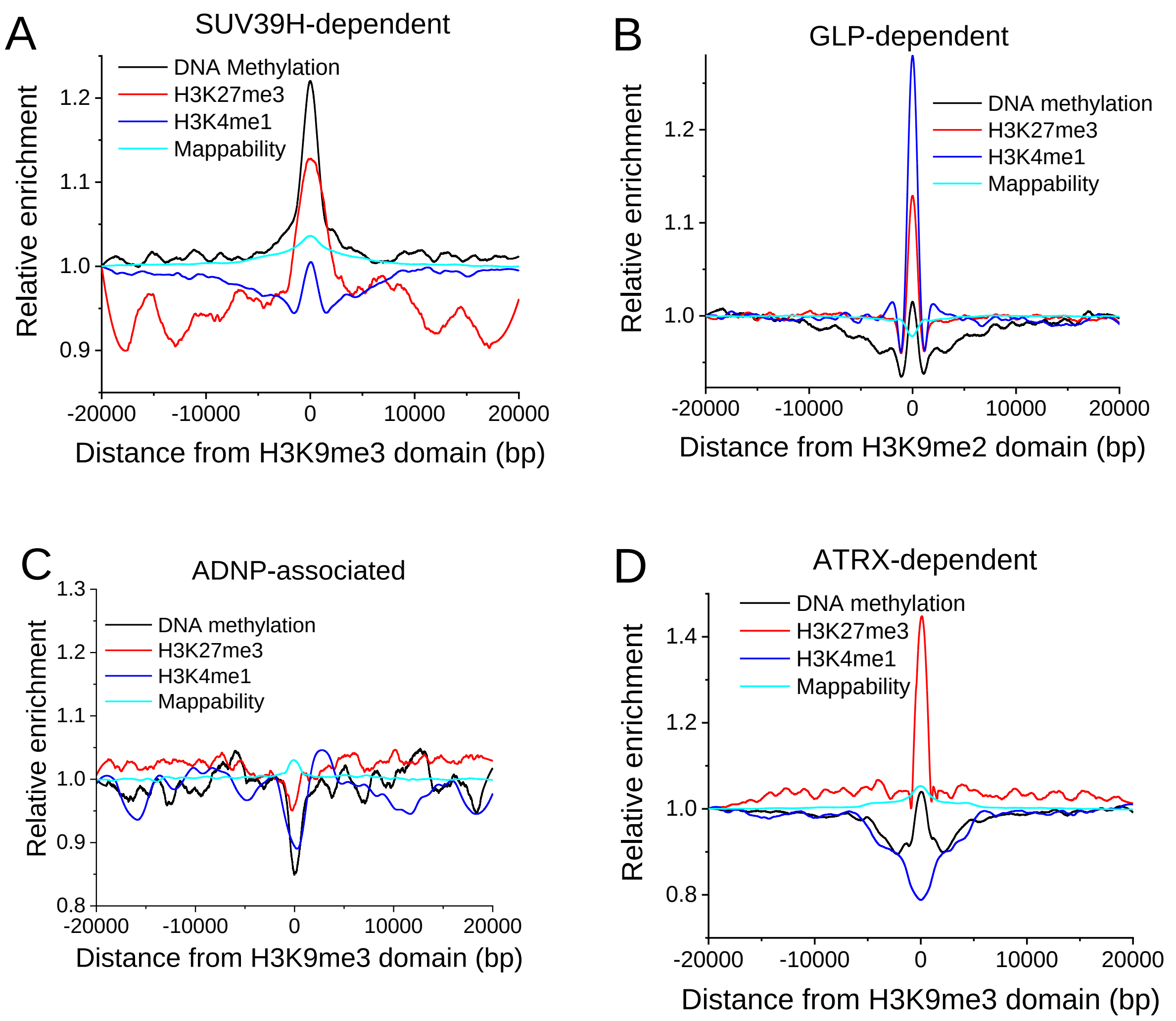


**Figure S1**. Average profiles of DNA methylation density, H3K27me3, H3K4me1 and mappability across the four types of heterochromatin regions shown in this manuscript: A) Suv39h dependent HNDs; B) GLP-dependent HNDs; C) ADNP-associated HNDs; D) ATRX-dependent HNDs. Note than, in contrast to the other three sets of data, mappability for the GLP-dependent peaks falls in the centre, indicating a higher proportion of repeating genomic elements in this dataset.

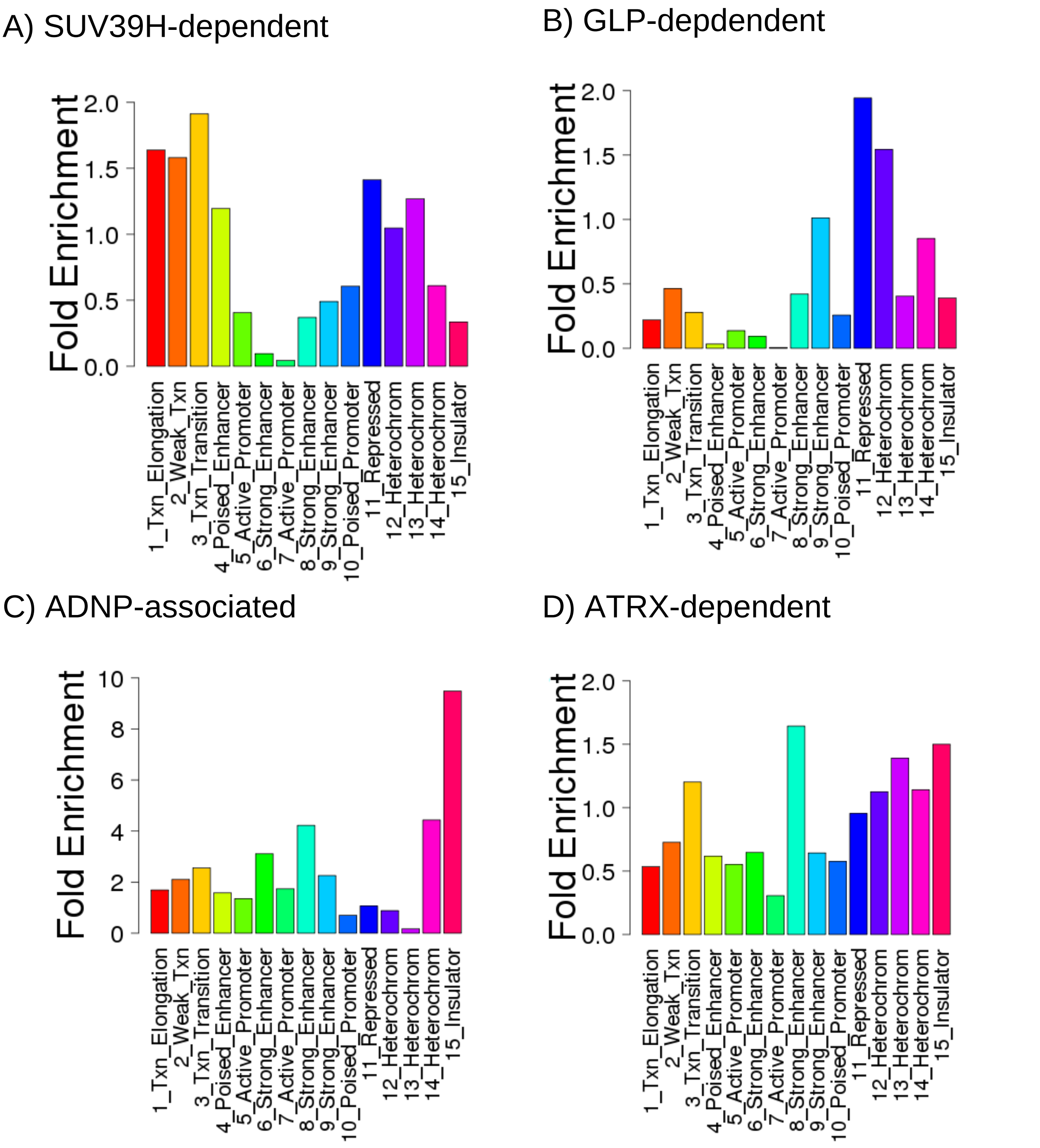


**Figure S2.** Enrichment of ChromHMM-determined ESC states [20] in the sets of heterochromatin peaks used in this manuscript. A) Suv39h-dependent HNDs. B) GLP-dependent HNDs. C) ADNP-associated HNDs. D) ATRX-dependent HNDs.

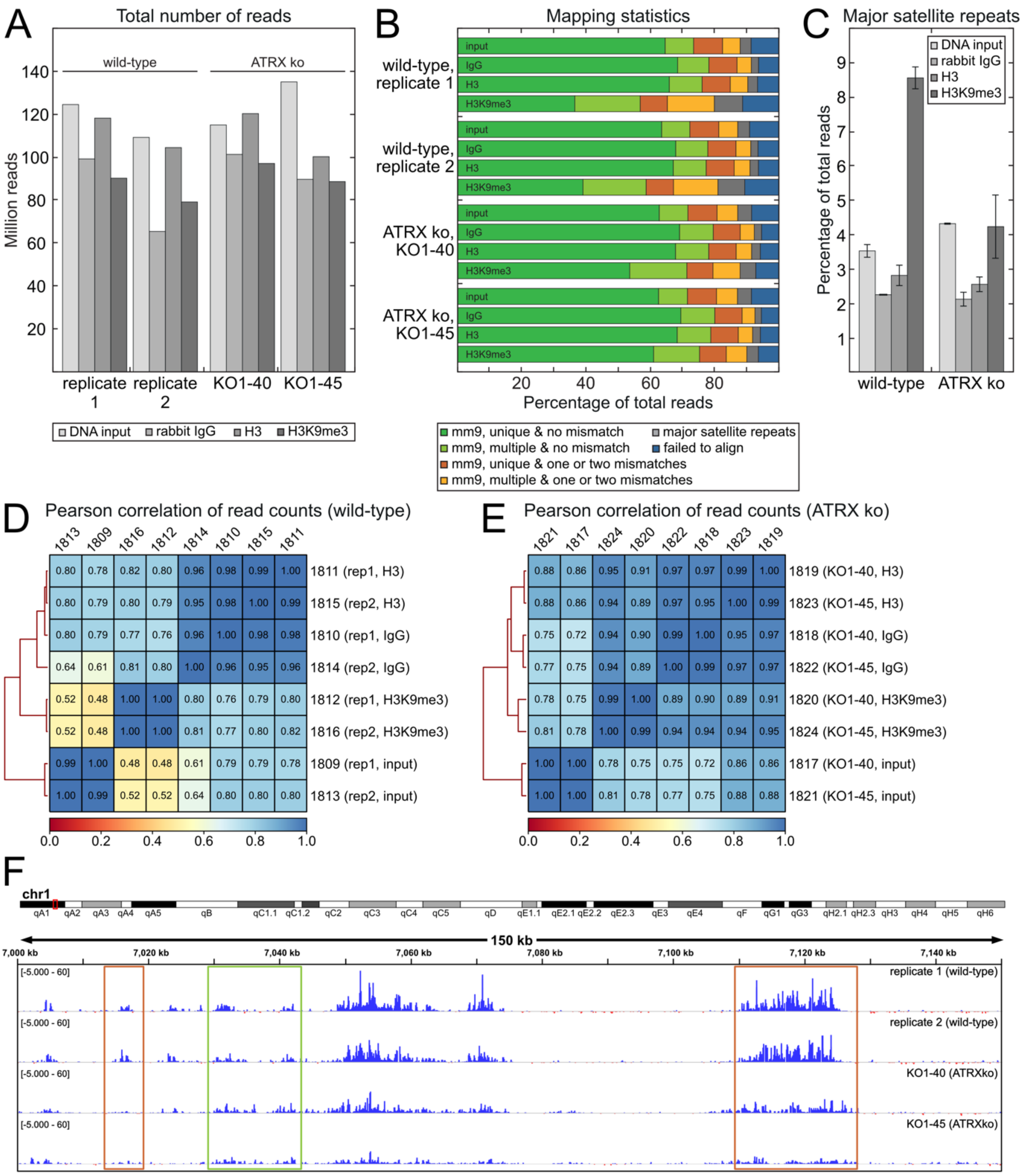


**Figure S3.** Quality control and basic mapping features of the H3K9me3 ChIP-Seq data set for wild-type and ATRX knockout mESCs. (**A**) Total number of reads from sequencing on the Illumina HiSeq 2000 platform. (**B**) Basic mapping statistics using Bowtie2 [21]. Notably, reads from H3K9me3 IP tend to map to multiple sites in the genome (yellow, light green) as expected since repetitive sequences are known to frequently carry H3K9me3 [22]. (**C**) Reads mapping the consensus sequence of major satellite repeats [23], a known H3K9me3 domain [24]. Notably, H3K9 trimethylation was reduced in ATRX ko mESCs. (**D**) Pearson correlation between read coverage of 10 kb bins throughout the genome calculated using deepTools2 [25] for wild-type samples. Importantly, replicates appear very similar to each other and the H3K9me3 IPs were rather dissimilar from the input. (**E**) Same a panel D but for ATRX ko samples. These samples indicating showed a lower enrichment over input, which is likely to reflect some loss of H3K9me3. (**F**) IGV traces [26] of normalized and background-corrected H3K9me3 traces for both wild-type and ATRX ko mESCs. Some regions appear rather independent of ATRX (green) whereas others showed less H3K9me3 enrichment in the ATRX ko samples compared to the wild-type (orange). Normalization and background correction were done as described previously using MCORE [27].

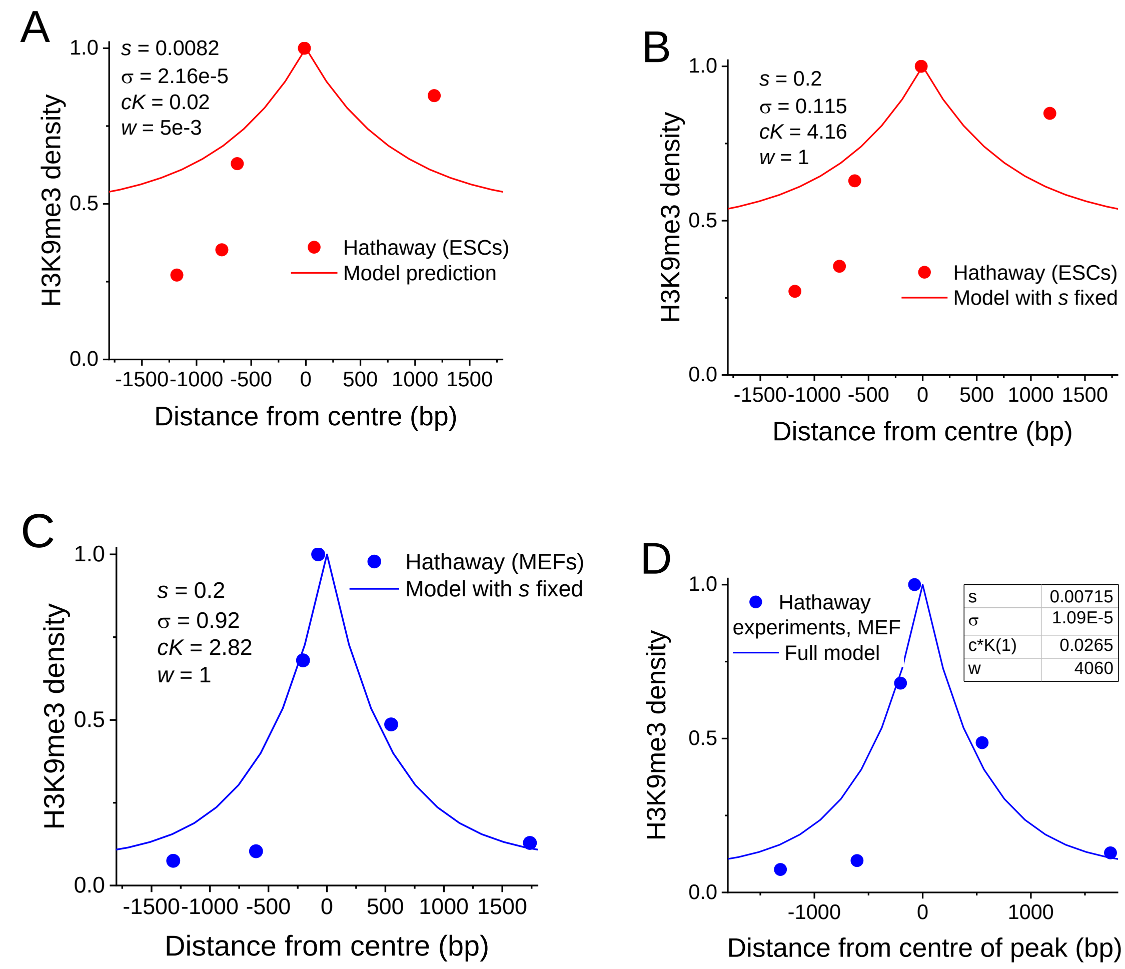


E

| | $s$ | $\sigma$ | $cK_1$ | $W$ |
|---|---|---|---|---|
| mESC | $8.18 \times 10^{-3}$ | $2.16 \times 10^{-5}$ | $2.02 \times 10^{-2}$ | $5.03 \times 10^{3}$ |
| MEF | $7.15 \times 10^{-3}$ | $1.09 \times 10^{-5}$ | $2.64 \times 10^{-2}$ | $4.06 \times 10^{3}$ |
| Suv39h-dep | $1.04 \times 10^{-2}$ | $1.46 \times 10^{-1}$ | $1.61 \times 10^{-2}$ | $5.11 \times 10^{3}$ |
| Atrx-dep | $4.29 \times 10^{-1}$ | $7.18 \times 10^{-6}$ | $2.55 \times 10^{-2}$ | $6.93 \times 10^{2}$ |
| GLP-dep | $7.10 \times 10^{-3}$ | $3.79 \times 10^{-9}$ | $2.37 \times 10^{-2}$ | $4.43 \times 10^{3}$ |
| mESC (*s* fixed) | 0.2 | $1.15 \times 10^{-1}$ | 4.16 | 1.00 |
| MEF (*s* fixed) | 0.2 | $9.19 \times 10^{-2}$ | 2.82 | 1.00 |

**Figure S4.** Comparison of predicted heterochromatin profiles with those determined in artificial heterochromatin experiments of Hathaway et al (2012). A) Artificially-induced heterochromatin in ESCs from Hathaway et al (2012) (full parameter search). B) The same as (A) but with fixed *s* parameter. C) Artificially-induced heterochromatin in MEFs from Hathaway et al (2012) (fixed *s* parameter, cf. panel (A). D) Artificially-induced heterochromatin in MEFs from Hathaway et al (2012) (full model). E) Table of best fit parameters *s*, $\sigma$ and w for the mESC and MEF data from Hathaway et al (2012), as well as the averaged profiles for Suv39h-dependent, ATRX-dependent and GLP-dependent HNDs in ESCs.

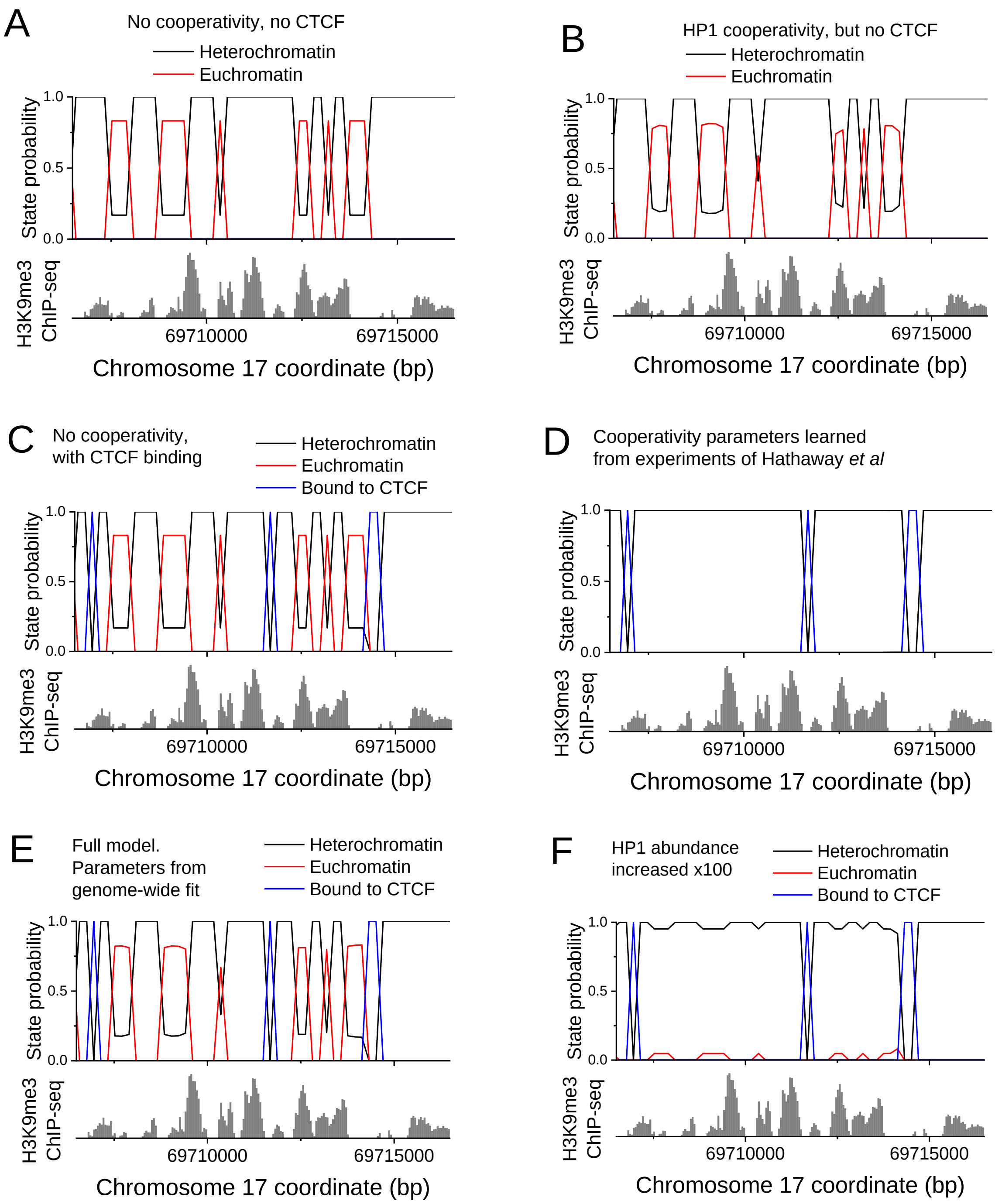


**Figure S5.** Model predictions (top panels) and experimental H3K9me3 ChIP-Seq profiles (bottom panels) for an example region containing Suv39h-dependent HNDs. A) No cooperativity of HP1 binding and no CTCF binding; B) HP1 binding cooperativity but no CTCF binding; C) CTCF binding but no HP1 cooperativity; D) HP1 cooperativity, CTCF binding and nucleosome-nucleosome interactions ($\sigma$ = 2.16e-5) learned from experiments of Hathaway *et al*); E) Full model fitted to genome-wide *in vivo* ChIP-seq data in ESCs. F) Same as (E) but the local HP1 concentration is increased 100-fold.

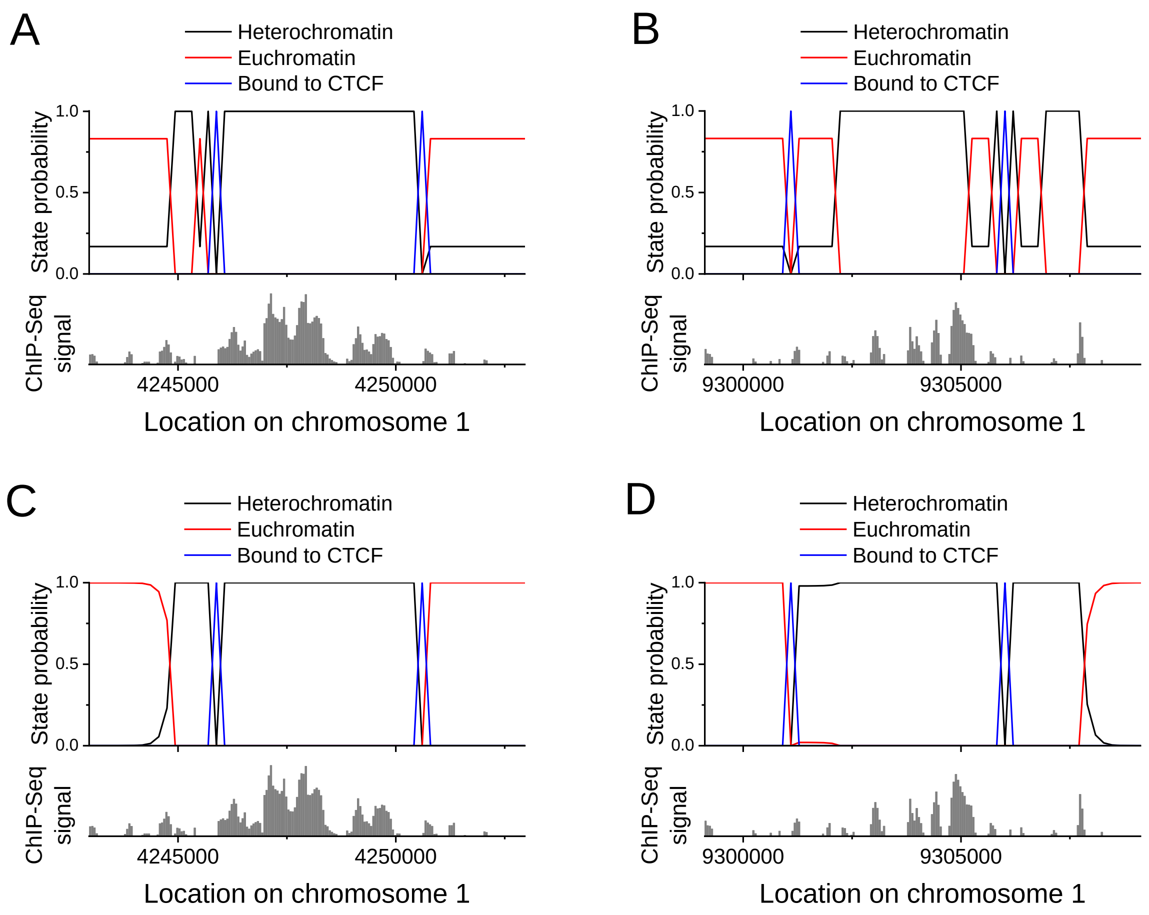


**Figure S6.** The effect of CTCF removal exemplified for two regions containing Suv39-dependent HNDs, chr1:4242961-4252961 and chr1:9299120-9309120. A and B) Predictions of the model using the best fit parameters from the Suv39-dependent heterochromatin peak size distribution, but without HP1-HP1 cooperativity. C and D) predictions of the model using the parameters fitted to the Hathaway et al artificially established heterochromatin in MEF. The effect of HP1-HP1 cooperativity results in smoothing the heterochromatin propagation front and expanding the nanodomain size.

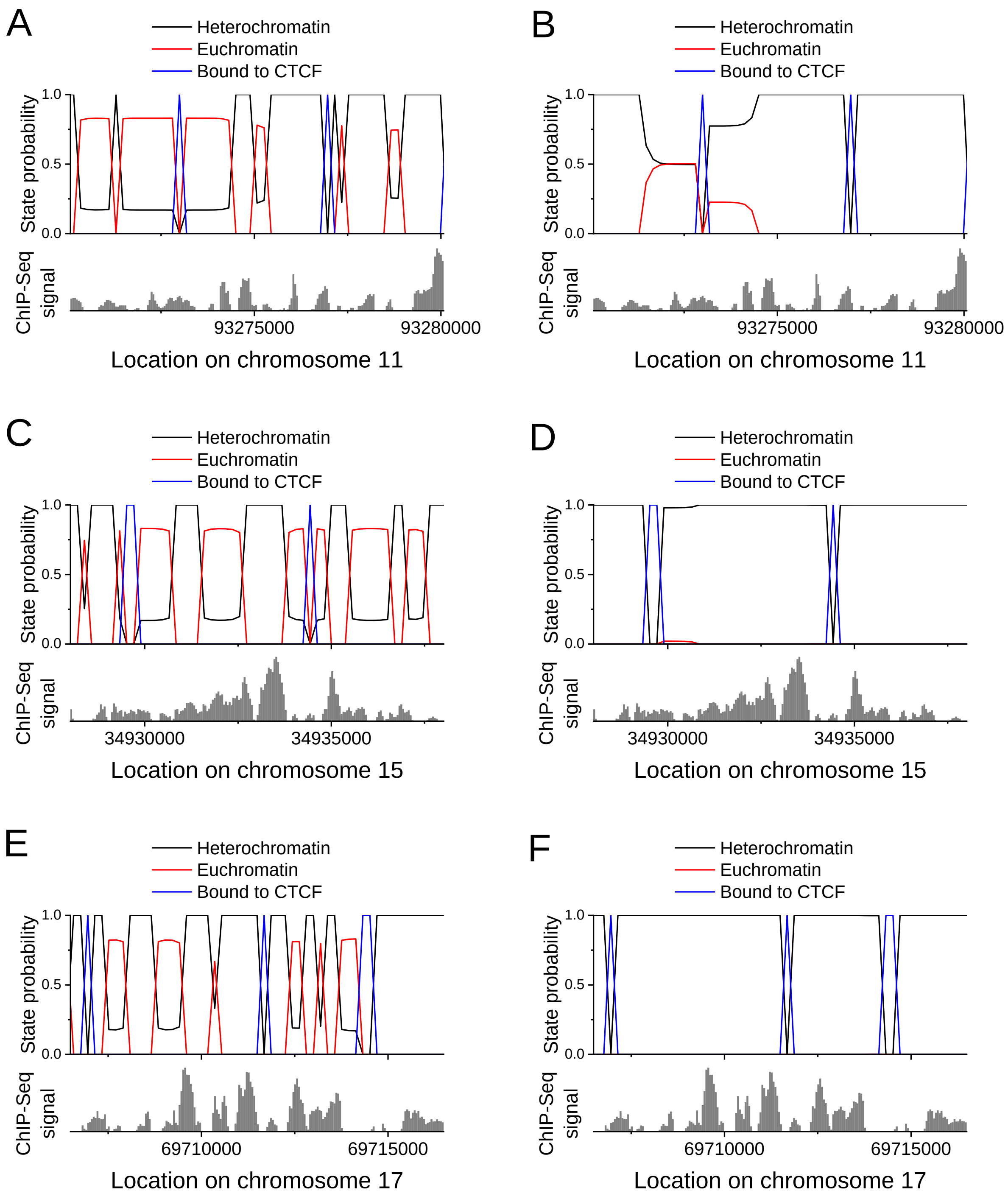


**Figure S7.** The effect of reducing $\sigma$ exemplified for three genomic regions containing several Suv39-dependent heterochromatin domains. Left-hand panels have been calculated for $\sigma = 1$; right-hand panels for $\sigma = 2.16 \times 10^{-5}$.

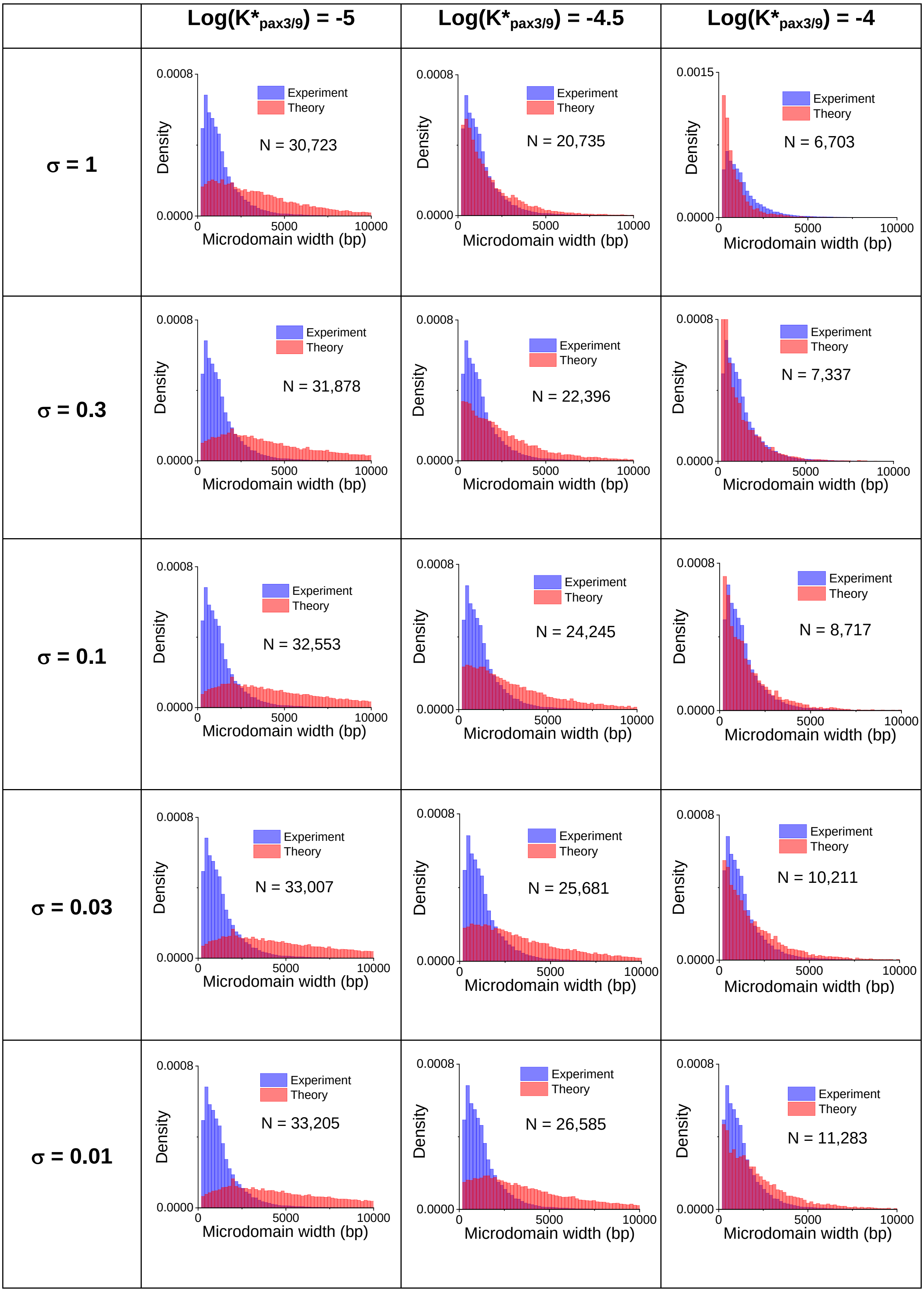

Log(K*pax3/9) = -5
Log(K*pax3/9) = -4.5
Log(K*pax3/9) = -4
σ = 1
σ = 0.3
σ = 0.1
σ = 0.03
σ = 0.01
Experiment
Theory
Density
Microdomain width (bp)
0.0008
0.0015
0.0000
0
5000
10000
N = 30,723
N = 20,735
N = 6,703
N = 31,878
N = 22,396
N = 7,337
N = 32,553
N = 24,245
N = 8,717
N = 33,007
N = 25,681
N = 10,211
N = 33,205
N = 26,585
N = 11,283

**Figure S8A.** Model predictions of Suv39h1/2-dependent nanodomain size distribution for varying values of the heterochromatin initiation threshold (Log($K^*_{pax3/9}$)**)** and σ (keeping other values fixed). Increasing either σ or Log($K^*_{pax3/9}$) reduces the average nanodomain size and the number of nanodomains containing initiation sites. The best fit is in the top-middle cell.

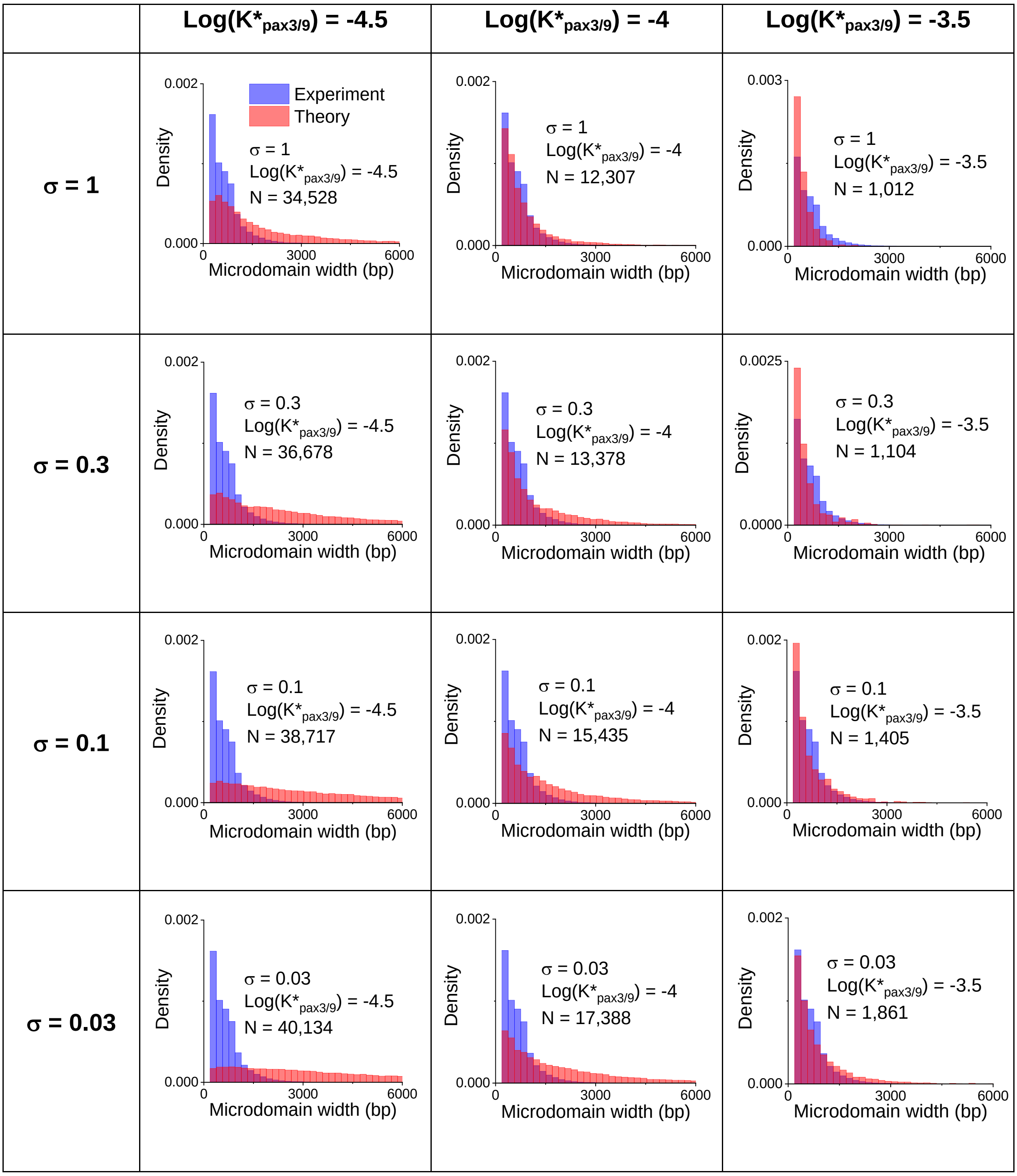

| | | | |
|---|---|---|---|
| **σ = 0.01** | 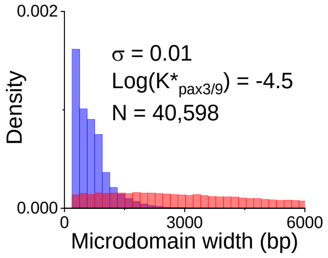 | 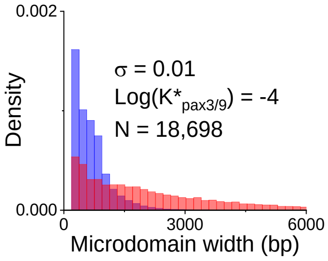 | 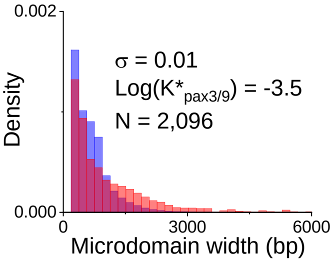 |







**Figure S8B.** Model predictions of GLP-dependent nanodomain size distribution for varying values of the heterochromatin initiation threshold (Log($K^*_{pax3/9}$)**)** and σ (keeping other values fixed). Increasing both σ and Log($K^*_{pax3/9}$) reduces the average size of nanodomains and the number of regions containing heterochromatin initiation sites. The parameters chosen for the main text are in the top-middle cell of this table.

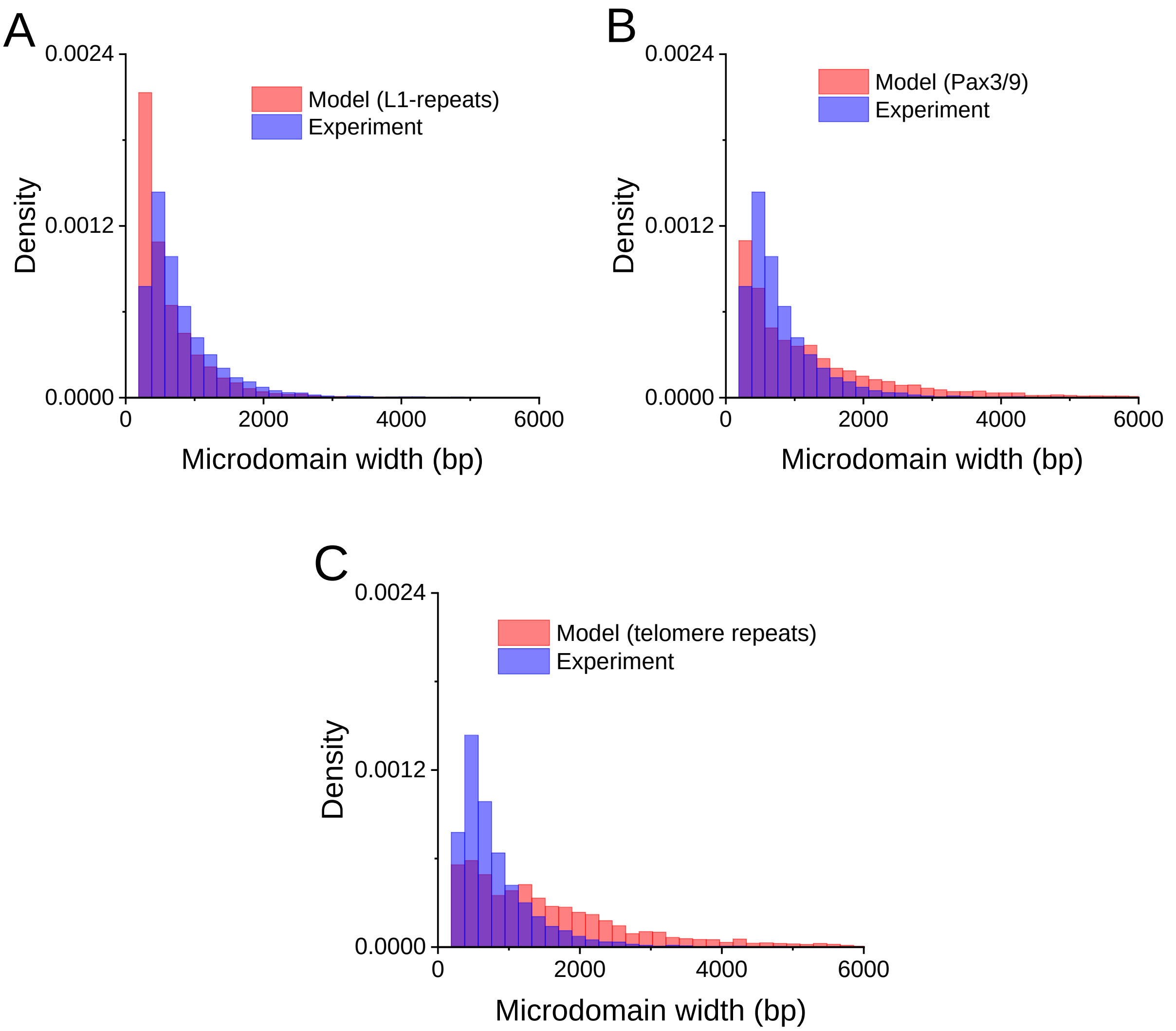


**Figure S8C.** Histograms of predicted peak distributions for the ATRX-dependent heterochromatin model using as initiation sites L1 repeats (A), PAX3/9 binding sites (B) and TERRA lncRNA-enriching telomere repeats (C).

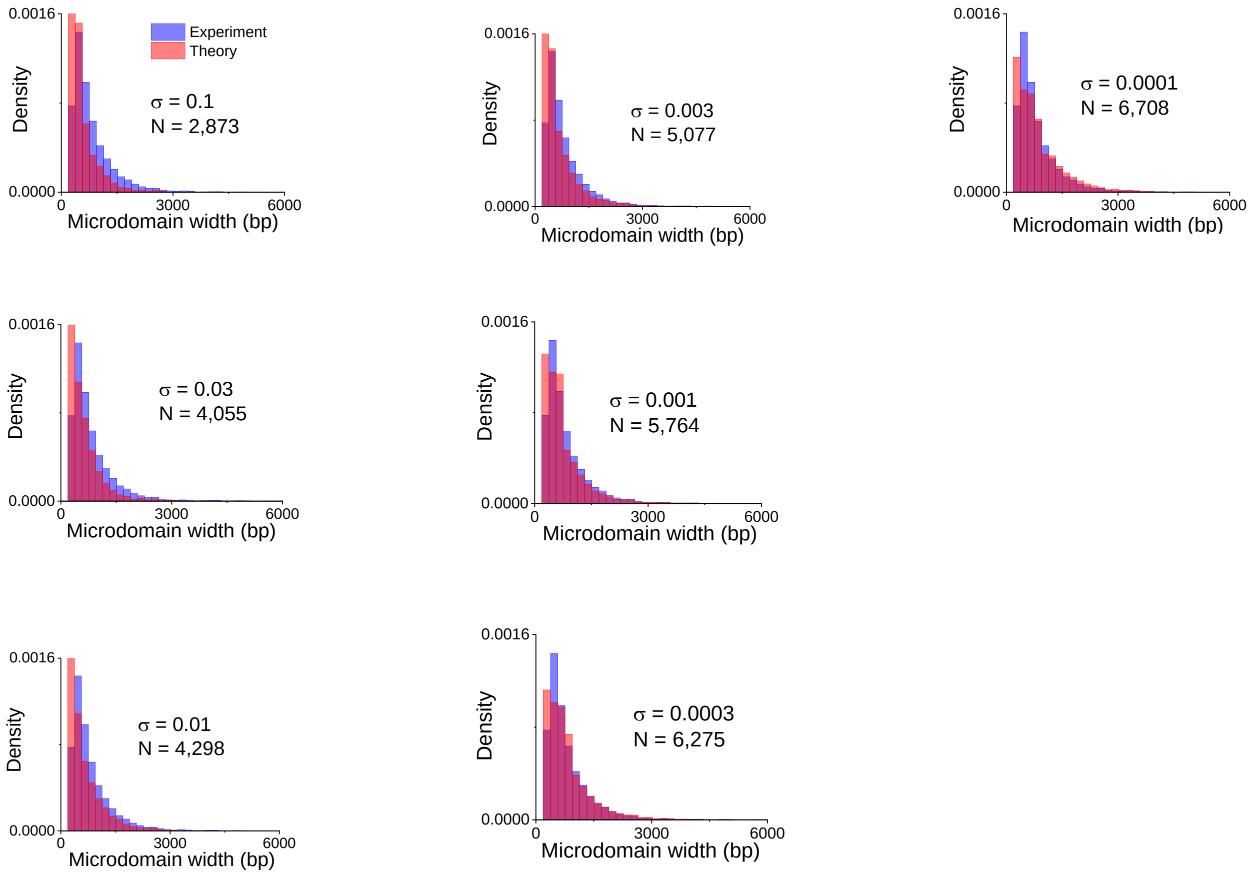


**Figure S8D:** Model predictions of ATRX-dependent nanodomain size distribution with experimental data for different choices of the boundary formation parameter $\sigma$ using the L1 repeat-initiating model. Increasing $\sigma$ reduces the size and number of peaks in the resulting distribution. In this case, heterochromatin is initiated in the model by the recognition of the repeat location, so the log-affinity threshold as it was in the Pax3/9-initiating model is not relevant. The choices of $\sigma$ are indicated on the figure.

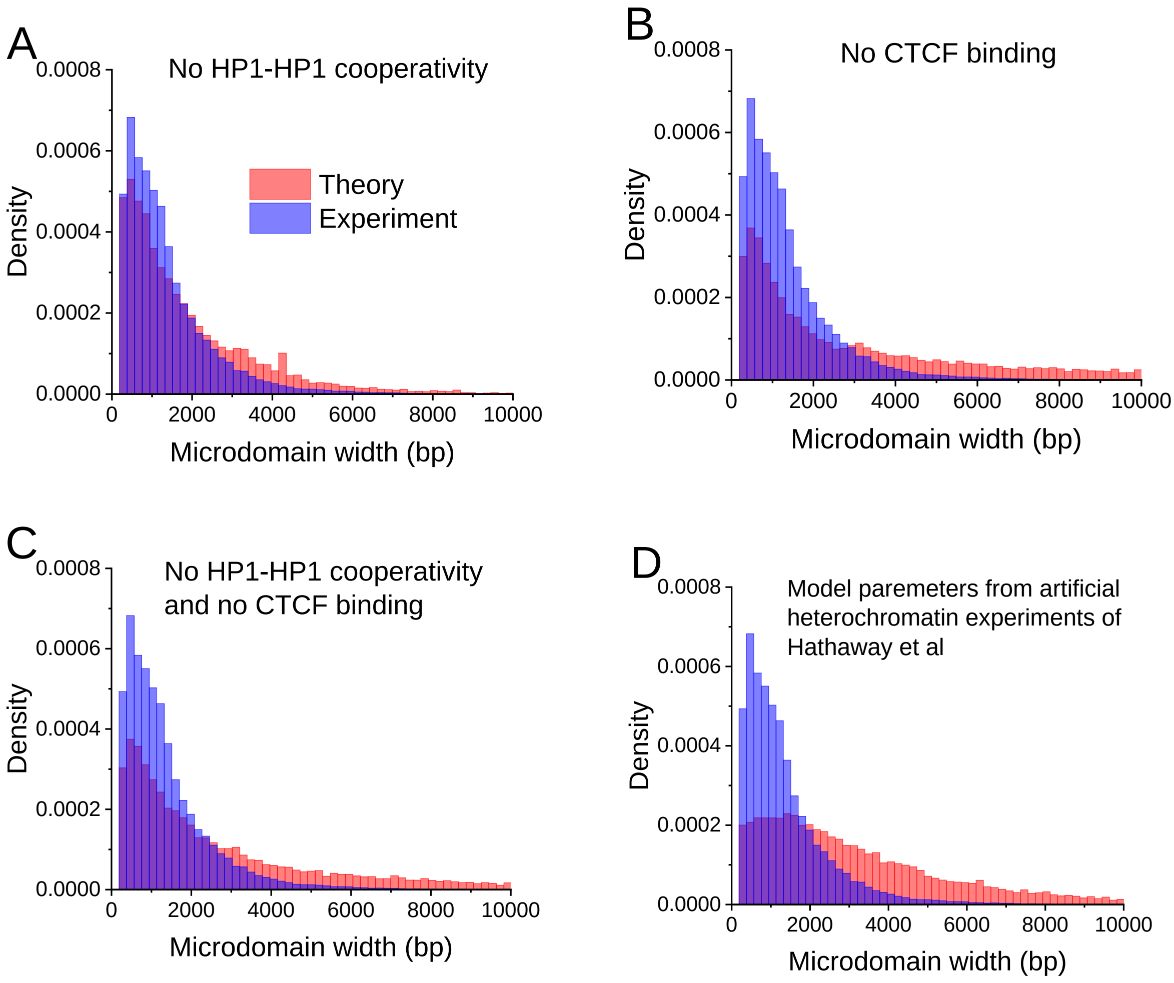


**Figure S9.** The effects of HP1-HP1 cooperativity and CTCF binding on Suv39h-dependent heterochromatin formation. (A) Model without HP1-HP1 cooperativity; (B) Model without CTCF binding; (C) Model without CTCF binding and HP1-HP1 cooperativity; (D) Model using the parameters derived from the Hathaway et al [10] experiments for artificial heterochromatin establishment in mESCs.

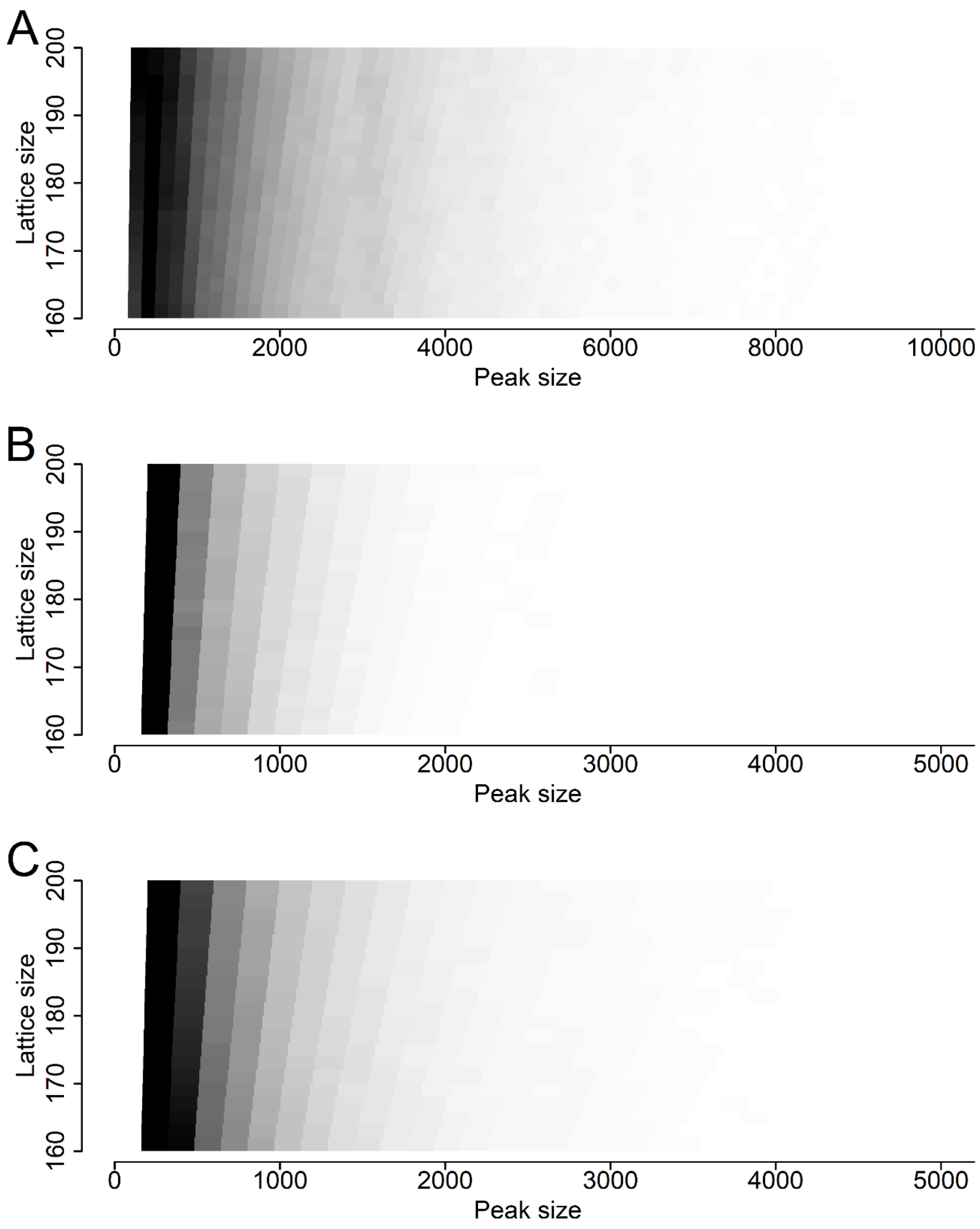
A
B
C
Lattice size
160
170
180
190
200
Peak size
0
2000
4000
6000
8000
10000
1000
3000
5000

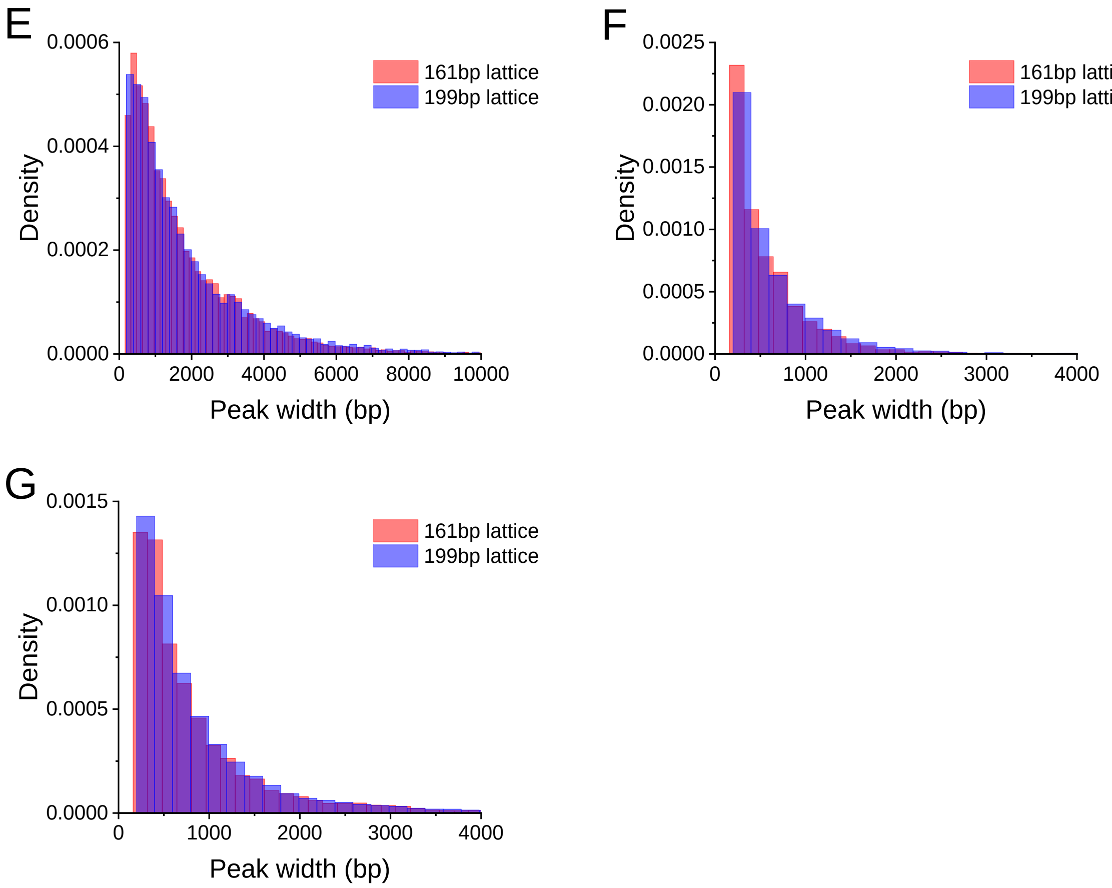


**Figure S10.** The effects in the assumed chromatin lattice unit size (NRL) on the nanodomain size distribution for Suv39h-dependent heterochromatin (A), ATRX-dependent heterochromatin (B) and GLP-dependent heterochromatin (C). Darker areas indicate more peaks of that size in the distribution. (E), (F) and (G) Comparisons of the predicted peak distributions for 161bp and 199bp lattice size for the (E) Suv39h, (F) ATRX-dependent and (G) GLP-dependent HNDs.

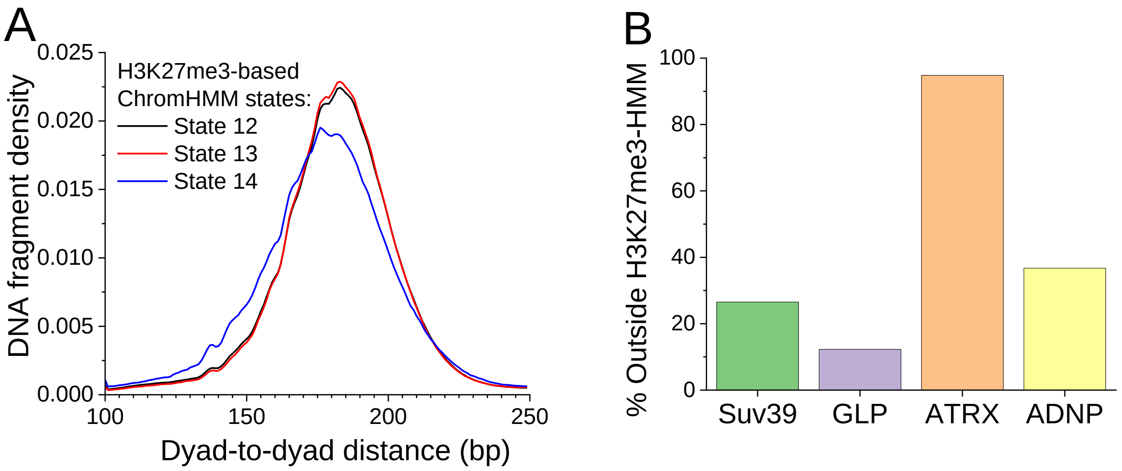


**Figure S11.** A) Nucleosome dyad-to-dyad distance distribution based on chemical mapping [28] for heterochromatin states determined using ChromHMM [20]. B) Nucleosome dyad-to-dyad distance distributions for Suv39-, ATRX-, GLP-dependent HNDs that do not intersect with any of the three ChromHMM heterochromatin states.

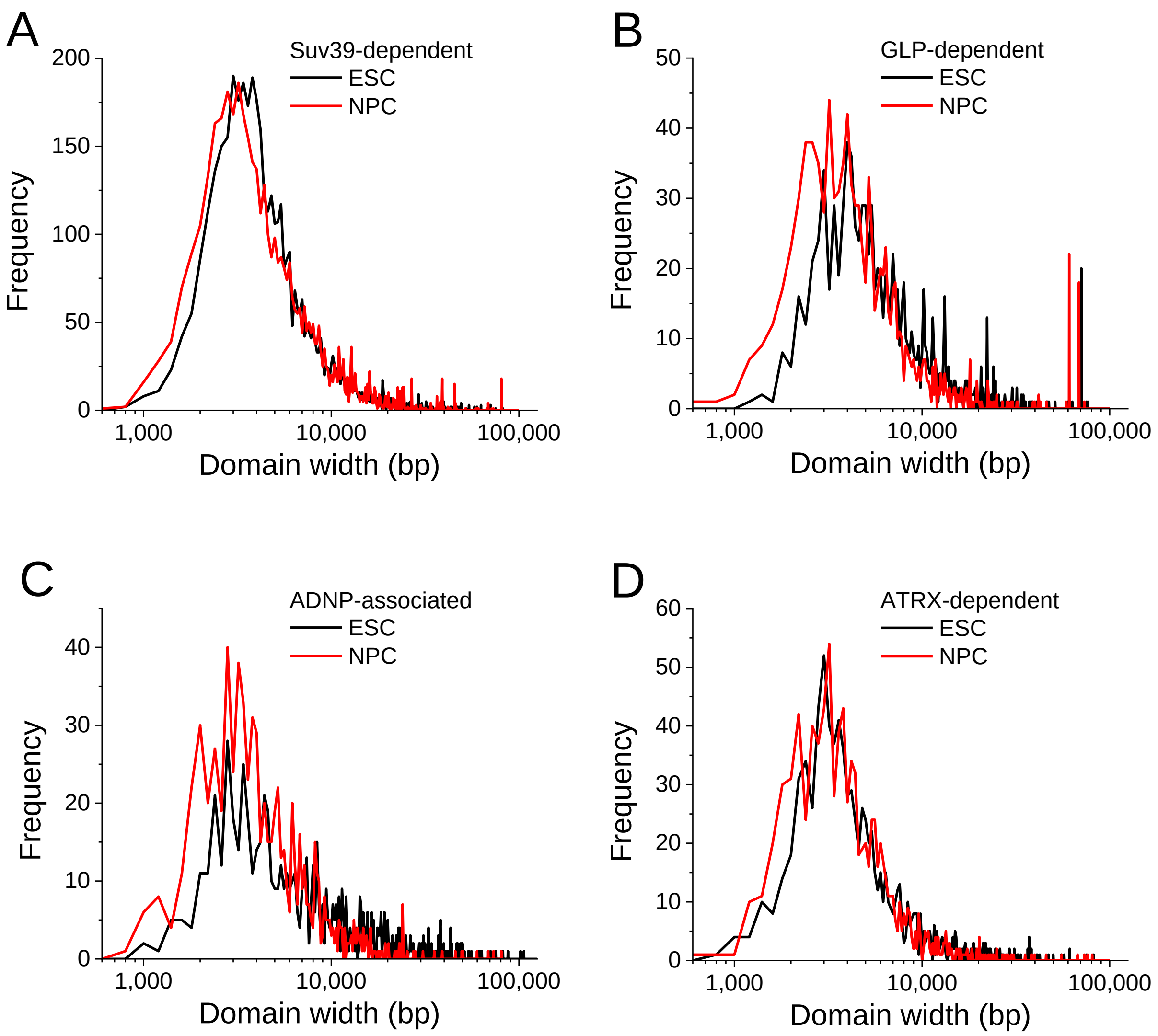


**Figure S12.** Experimental distribution of sizes of H3K9-methylated domains in ESCs in comparison with Neural Progenitor Cells (NPCs) differentiated from them, for different types of heterochromatin defined in ESCs. Only heterochromatin regions which overlap between ESCs and NPCs are considered. A) Suv39-dependent (N=4510), B) GLP-dependent (N=1055), C) ADNP-associated (N = 799), D) ATRX-dependent H3K9-methylated domains (N = 961).

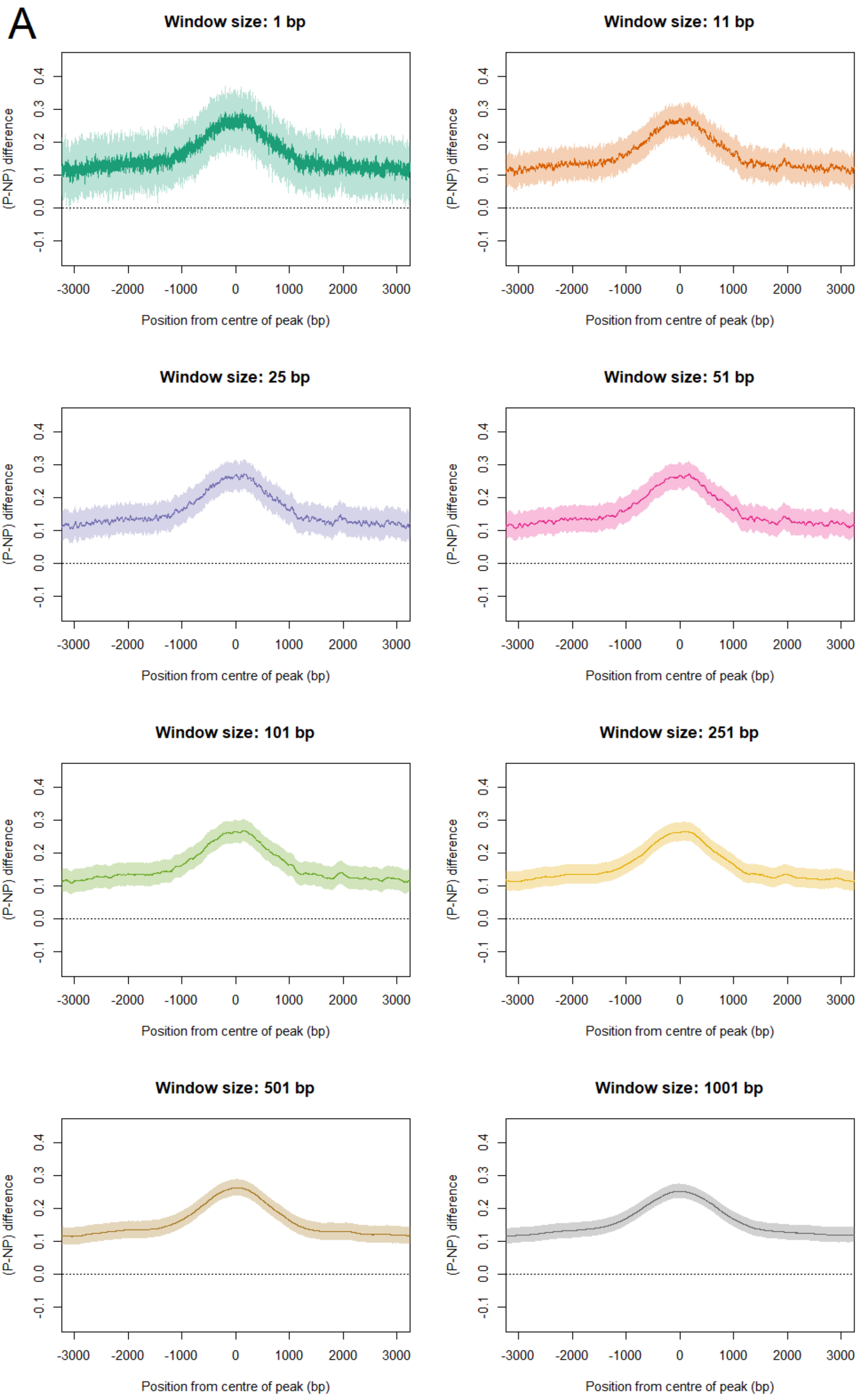
A
Window size: 1 bp
Window size: 11 bp
Window size: 25 bp
Window size: 51 bp
Window size: 101 bp
Window size: 251 bp
Window size: 501 bp
Window size: 1001 bp
(P-NP) difference
-0.1 0.0 0.1 0.2 0.3 0.4
-3000 -2000 -1000 0 1000 2000 3000
Position from centre of peak (bp)

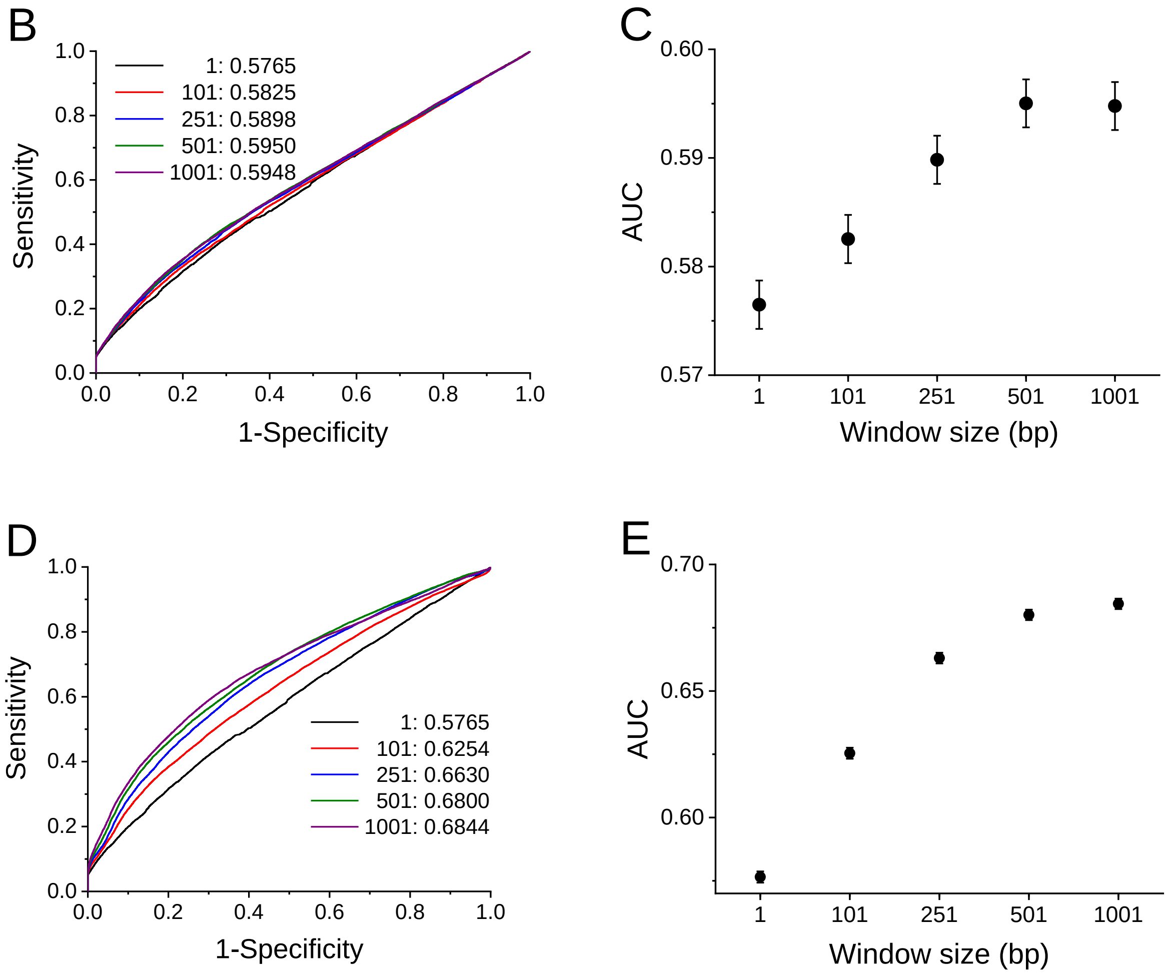


**Figure S13.** TRAP affinity scores calculated across Suv39h-dependent heterochromatin regions for different values of smoothing window: 1, 101, 251, 501 and 1001-bp centred geometrically-averaged windows. A) The difference between the log affinity profile geometrically averaged across all peaks and the profile geometrically averaged across non-peak regions: a ± 5 standard error is shown either side of the average. B) ROC curves based on the TRAP score with the matched non-peak regions, with AUCs for each window size shown. C) AUCs for each window size with estimated standard error. (D) and (E) ROC and AUC values using arithmetical-smoothing for the same conditions as (B) and (C).

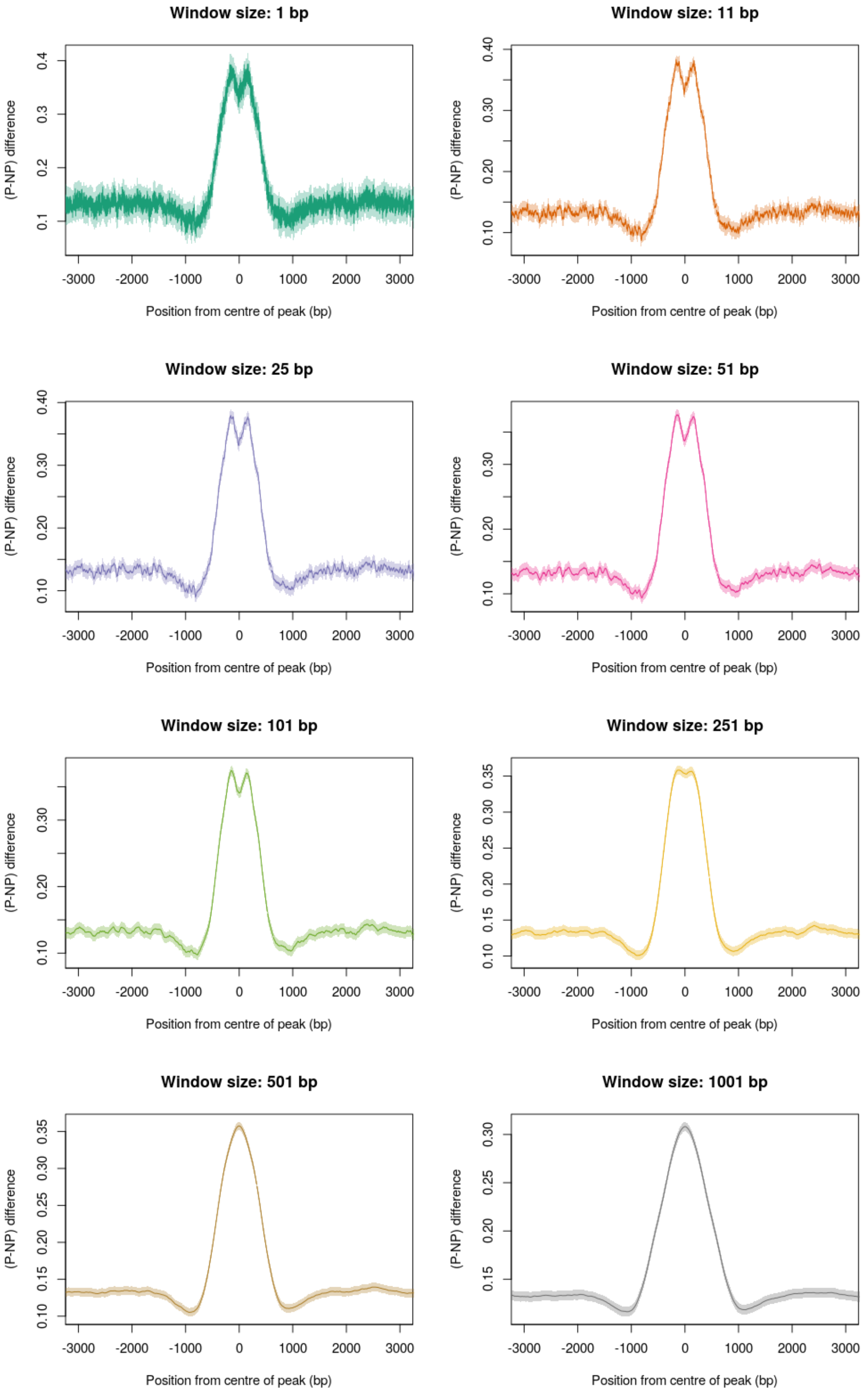


**Figure S14.** TRAP affinity scores calculated across GLP-dependent heterochromatin for different values of smoothing window, 1, 101, 251, 501 and 1001-bp centred geometrically-averaged windows, difference between the log affinity profile geometrically averaged across all peaks and the profile geometrically averaged across non-peak regions: a $\pm$ 2 standard error is shown either side of the average.

| Repeat name | | Repeat class | | Repeat family | |
|---|---|---|---|---|---|
| 686 | L1Md_F2 | 9483 | LTR | 5643 | L1 |
| 668 | L1Md_T | 5853 | LINE | 4479 | 4ERVK |
| 472 | B3 | 3550 | SINE | 3002 | MaLR |
| 432 | L1Md_A | 2335 | Simple_repeat | 2335 | Simple_repeat |
| 408 | MERVL-int | 421 | Low_complexity | 1172 | ERV1 |
| 335 | ORR1D1 | 405 | DNA | 1075 | Alu |
| 321 | B3A | 139 | Satellite | 1054 | B2 |
| 318 | RLTR10 | 80 | Other | 980 | B4 |
| 315 | B4A | 46 | Unknown | 827 | ERVL |
| 285 | L1Md_F | 34 | rRNA | 420 | Low_complexity |
| 275 | Lx8 | 11 | scRNA | 313 | MIR |
| 249 | ID_B1 | 10 | snRNA | 258 | MER1_type |
| 247 | L1_Mus1 | 9 | tRNA | 173 | L2 |
| 233 | RSINE1 | 4 | RNA | 138 | Satellite |
| 231 | (CA)n | 2 | srpRNA | 124 | ID |
| 220 | MTD | 1 | RC | 80 | Other |
| 216 | (TG)n | | | 78 | MER2_type |
| 209 | ETnERV2-int | | | 42 | Unknown |
| 185 | (TCTA)n | | | 34 | rRNA |
| 183 | B2_Mm2 | | | 32 | CR1 |

**Table S1.** Most frequent repeats overlapping ATRX-dependent heterochromatin peaks as reported by the RepeatMasker track for *Mus musculus* (mm9) on the UCSC Genome Browser, ordered by the repeat name as given, the class of repeat and the family of repeat.
